# Lipid-mediated GPR32 signaling reprograms macrophage metabolism to impair anti-tuberculous immunity

**DOI:** 10.1101/2025.07.07.663524

**Authors:** Joaquina Barros, Leandro Ferrini, Mariano Maio, Martín Manuel Ledesma, María Emilia Boix, Natacha Faivre, Sarah C. Monard, José Luis Marín Franco, Arnaud Métais, Florencia Sabbione, Federico Fuentes, Tomas Martín Grosso, Agustina Juliana Errea, Xavier Aragone, Milagro Sanchez Cunto, Domingo Palmero, Mario Matteo, Matías Ostrowski, Rafael J Argüello, Ranjan Kumar Nanda, Emilie Layre, Olivier Neyrolles, Christel Vérollet, Geanncarlo Lugo Villarino, Luciana Balboa

## Abstract

Mycobacterium tuberculosis, the causative agent of tuberculosis (TB), has evolved strategies to evade innate immunity and establish persistent infection. However, the mechanisms by which M. tuberculosis reprograms human macrophage metabolism remain incompletely defined. Tuberculous pleural effusion (TB-PE), a common extrapulmonary manifestation that frequently coexists with pulmonary TB, offers a unique, clinically relevant immunometabolic window into the TB microenvironment. Here, using patient-derived TB-PE samples, we demonstrate that this microenvironment induces a metabolic state in human macrophages that compromises their antimicrobial function. Lipidomic analysis identified an enrichment of the specialized pro-resolving mediator Resolvin D5 (RvD5), which signals through GPR32 to suppress macrophage microbicidal activity. The acellular fraction of TB-PE was sufficient to induce RvD5 secretion by monocytes, correlating with increased expression of RvD5 biosynthetic enzymes in pleural monocytes from TB patients. Mechanistically, RvD5-GPR32 signaling inhibited glycolysis without promoting oxidative phosphorylation, reducing HIF-1α activity and impairing intracellular M. tuberculosis control. HIF-1α stabilization restored antimicrobial function. These findings uncover the RvD5-GPR32-HIF-1α axis as a mechanism of metabolic immune suppression and a potential target for host-directed TB therapy.

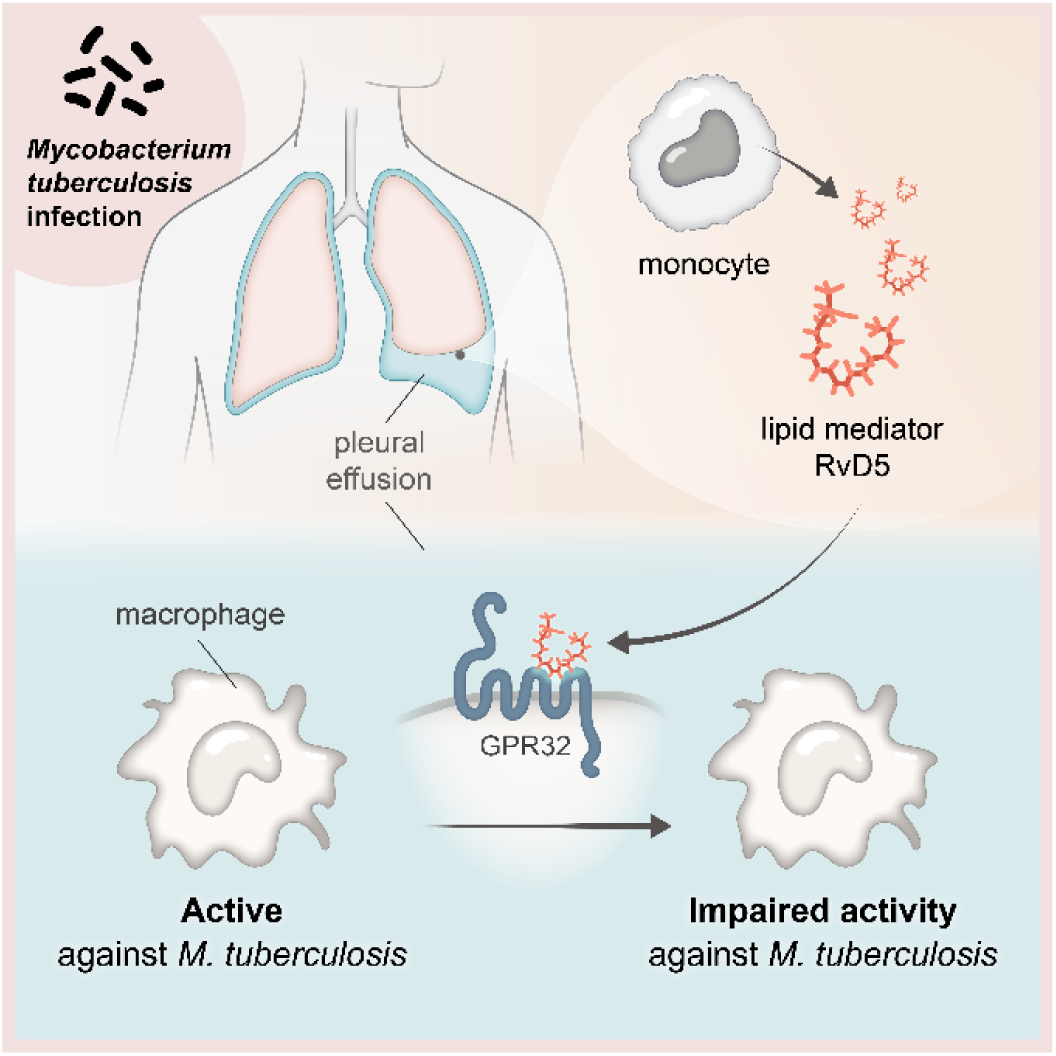

## Introduction

The success of *M. tuberculosis* (Mtb) as a pathogen lies, in part, in its efficient adaptation to the intracellular environment of human macrophages, which involves regulating the host’s metabolic pathways to facilitate immune evasion. Macrophages are essential immune cells that can serve as a niche for intracellular mycobacterial persistence but can also limit infection by activating their microbicidal properties. In fact, innate immunity appears sufficient to clear Mtb infections in certain patients, conferring resistance^1^. This activation of macrophages towards the microbicidal program (called the M1 activation program) depends on a profound metabolic change that promotes glycolysis to the detriment of mitochondrial oxidative phosphorylation^2,3^.

Tuberculous pleural effusion (TB-PE) is characterized by an accumulation of fluid in the pleural cavity of infected patients, marked by elevated levels of proteins, lipids, and specific leukocyte populations. Recent studies have begun to characterize the metabolites present in these samples, revealing differential abundance of amino acids, citric acid cycle intermediates, and free fatty acids compared with malignant pleural effusions^4^. TB-PE is typically a serous fluid and contains very few viable mycobacteria, indicating a paucibacillary infection. The development of TB-PE is likely due to a low-burden mycobacterial infection in the pleural space, which typically arises from initial parenchymal lesions. This infection triggers an immune response that not only increases the production of pleural fluid but also impairs its drainage^5^.

TB-PE is the second most prevalent form of extrapulmonary TB, after TB lymphadenitis, accounting for 15-25% of TB cases worldwide. In immunocompromised populations, it may represent as much as 50% of all cases^6^. Compared to samples obtained from blood or bronchoalveolar lavage (BAL) of TB patients, TB-PE provides a more accurate reflection of the microenvironment encountered during infection. It captures the specific cellular composition and environmental factors at the site of infection, which is anatomically related to the lungs. In fact, there is a significant association between pleural TB and pulmonary TB as coexisting parenchymal disease has been identified in 48%^7^ and 86%^8^ of computed tomography scans conducted on patients with pleural TB. Furthermore, TB-PE samples better reflect the physiological concentrations of metabolites and compounds because they are not diluted with saline, unlike BAL. These distinctive characteristics make TB-PE a valuable tool for TB research.

While it has been suggested that type-1 immunity is dominantly present in TB-PE^9^, infection control within the pleural cavity is considered insufficient, as patients frequently experience recurrent episodes of TB if left untreated^10^. This raises the possibility of active mechanisms that impede the development of a long-term protective immune response. In this regard, we demonstrated that the acellular fraction of TB-PE induces the differentiation of monocytes into immunosuppressive macrophages^11–13^, converting them into lipid-laden or foamy macrophages, thereby affecting their effector functions against bacilli^14^. We have also demonstrated that microbicidal M1 macrophages exposed to a lipid fraction isolated from TB-PE, enriched in metabolites derived from polyunsaturated fatty acids (PUFA), are unable to carry out the metabolic reprogramming associated with their pro-inflammatory profile, showing reduced glycolytic activity, as well as increased mitochondrial respiration and reduced microbicidal activity^15^; however, the mechanism of action as well as the specific metabolites involved in this effect remain unclear, limiting the potential to develop targeted therapies aimed at enhancing anti-TB immune response.

In response to infection, both inflammatory and pro-resolving processes are deployed in TB-PE. Among the mechanisms aimed at limiting the inflammatory process and activating tissue repair and healing are those catalyzed by a family of lipids, recently termed specialized pro-resolving mediators (SPMs)^16^. SPMs derive from PUFA such as linoleic acid (C18: Δ2, n-6), arachidonic acid (C20: Δ4, n-6), eicosapentaenoic acid (EPA, 20:5, n-3), and docosahexaenoic acid (DHA, 22:6, n-3), which are produced through enzymatic reactions catalyzed by lipoxygenases (LOX) and/or cyclooxygenases (COX). SPMs are composed of DHA-derived D-series protectins and resolvins, EPA-derived E-series resolvins, and arachidonic acid-derived lipoxins.

By binding to specific receptors, resolvins modulate various inflammatory processes, including neutrophil migration, macrophage phagocytosis, and the expression of pro-inflammatory mediators, thereby mitigating inflammatory pathologies^17^. The G protein-coupled receptor 32 (GPR32) has been shown to bind to resolvins RvD1, RvD3, and RvD5^18–20^. Although direct evidence supporting the RvD5-GPR32 axis in lung infections remains limited, investigations into related resolvins, such as RvD1^21^, yield promising results in models of lung injury, indicating reduced neutrophil infiltration and diminished alveolar damage. These findings strongly implicate the RvD1-GPR32 axis as a critical regulatory pathway. This mechanism is especially pertinent in the context of bacterial pneumonia and viral infections, like influenza, where a carefully calibrated immune response is essential. Despite the known importance of resolvins, no study to date has investigated the role of RvD5 in TB. In this study, we performed a lipidomic analysis of TB-PE and found high levels of the lipid mediator RvD5. We characterized its function by exposing M1 macrophages to this lipid mediator and found that RvD5 is sufficient to impair their microbicidal activity and glycolytic metabolism. We functionally validate that the direct binding mediates this impairment between RvD5 and its receptor GPR32, expressed in macrophages, whose activation inhibits HIF1α. Restoring HIF1α with a chemical activator was sufficient to rescue the functionality of M1 macrophages exposed to RvD5.

## Results

### Enrichment in specialized resolution lipid mediators correlates with impaired macrophage glycolysis in tuberculous pleural effusions

We previously found that PUFA metabolites in TB-PE inhibit the glycolytic activity of M1 macrophages^15^. To explore whether this inhibition was specific to pleural TB lipids, we compared the ability of the PUFA extracted from the acellular fraction of TB-PE and non-TB patients to inhibit glycolysis in M1 macrophages (activated by IFNγ and LPS), by assessing lactate release from these cells. The lipid fractions isolated from TB-PE showed greater inhibition of lactate release by M1 macrophages than PE from non-tuberculous patients with other pathologies, such as heart failure, pneumonia, and cancer (**Figure 1A**). To further investigate the candidate components responsible for this inhibition, a targeted lipidomic analysis was conducted on lipids isolated from TB and non-TB patients (n=8 and 3, respectively). Among the 29 screened PUFA metabolites, four exhibited higher levels in TB-PE compared to non-TB PE (**Figure 1B-C**). Notably, these molecules were specialized pro-resolving lipid mediators (SPMs) derived from omega-3 fatty acids, i.e., 14-HDoHE, RvD5, Protectin Dx (PDx), and 18-HEPE (**Figure 1B-C**). Importantly, the abundance of these SPMs was significantly correlated with the decline in glycolysis induced by each PE in monocyte-derived M1 macrophages from healthy donors (**Figure 1D**). Taken together, our data indicate that SPMs are highly abundant in TB-PE, and their levels correlate with TB-PE’s capacity to suppress M1 macrophage glycolytic activity. Moreover, RvD5 was significantly enriched in TB-PE compared to non-TB PE, with a log₂ fold change of 15.07, a−log(p-value) of 3.09, and a multiple testing-corrected q-value of 0.02. By contrast, 18-HEPE and 14-HDoHE showed only marginal statistical significance (q = 0.05) (**Figure 1C and E**).

**Figure 1.**
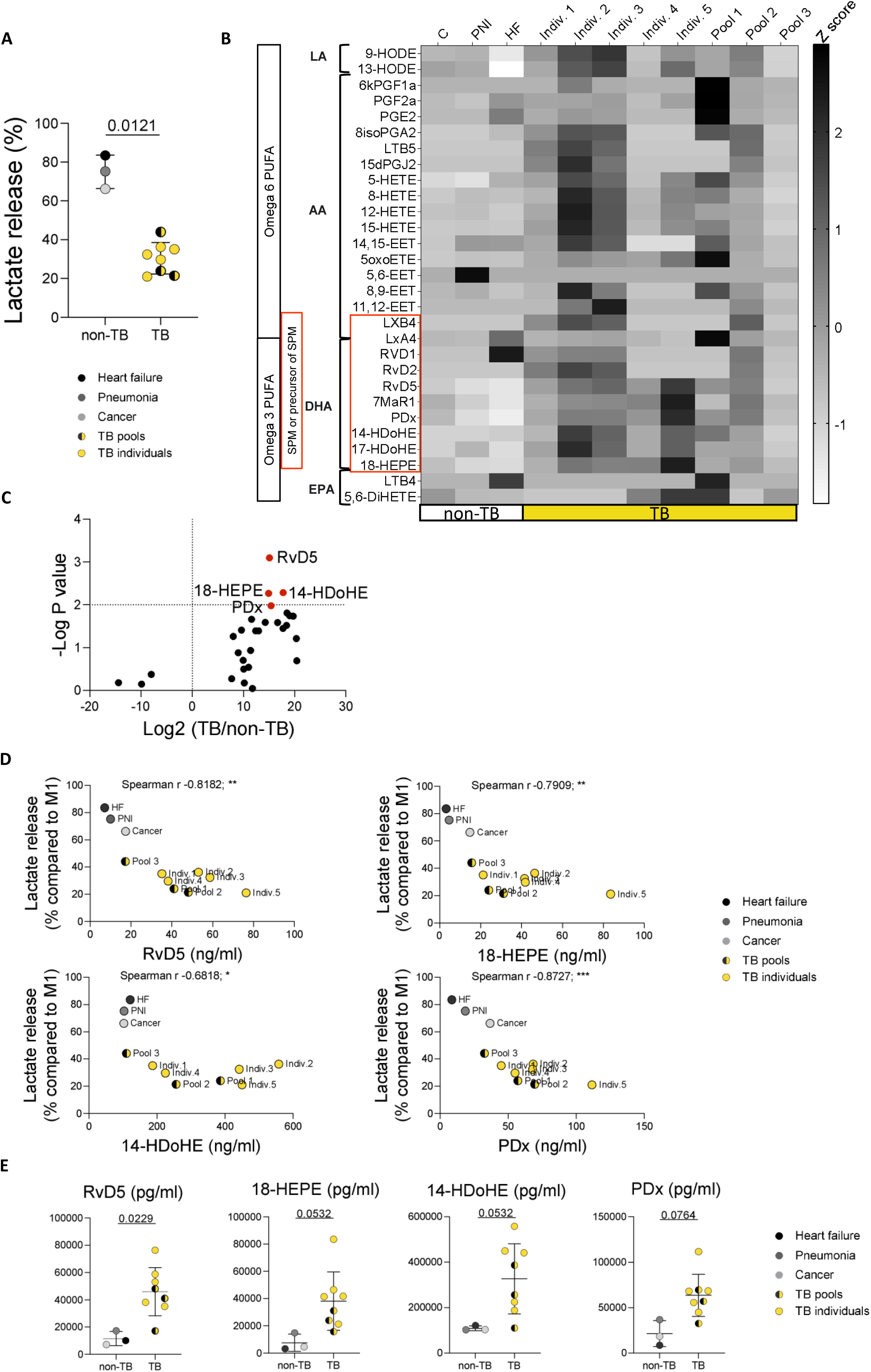
Enrichment in specialized resolution lipid mediators correlates with impaired macrophage glycolysis in tuberculous pleural effusions. (A) M1 macrophages were exposed or not to the polyunsaturated fatty acid (PUFA) extracted from the acellular fraction of tuberculous (TB; three pools and five individual samples) or non-tuberculous (non-TB; three pools) effusions. The percentage of release of lactate in the supernatant compared to untreated M1 cells is shown. Data are presented as individual values with the mean ± SD. Statistical significance was determined by a Mann-Whitney test. In the figure, sample pools are distinguished by color and symbol. Effusions from tuberculous (TB) patients are shown as yellow circles 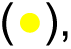 with pooled samples containing black dots. The three cardiac failure pools are black circles (●), the single pneumonia pool is dark gray (●) and the single malignancy pool is light gray represent the single malignancy pool 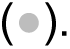 (B) Heat map illustrating the relative abundance (Z-score normalized) of 29 PUFA lipid species detected through lipidomic analysis in pleural effusions classified as TB and non-TB. Lipids are grouped according to their precursor PUFA. The specialized pro-resolving mediator (SPM) family is highlighted (red box). Non-TB effusions comprise heart failure (HF), pneumonia (PNI), and cancer (C). (C) The volcano plot depicts fold changes in lipid levels between TB (n=8) and non-TB (n=3) pleural effusion samples, along with their statistical significance (p-value), for all detected metabolites. (D) Correlations between lactate release by M1 macrophages exposed to TB or non-TB pleural effusions and the abundance of specific lipids. Samples and pool symbols are consistent with panel A. Spearman’s correlation coefficient (r) and significance levels are reported. (E) Abundances of selected SPMs (RvD5, 14-HDoHE, 18-HEPE, and PDx). Samples and pool symbols are consistent with panel A. A two-way ANOVA with Benjamini, Krieger, and Yekutieli correction for multiple comparisons was performed. Corrected P values (q value) are shown. LA: linoleic acid, AA: arachidonic acid, DHA: docosahexaenoic acid, EPA: eicosapentaenoic acid.

To ensure that the inclusion of pooled samples in the lipidomic analysis did not bias these results, we validated that pooled TB-PE samples behave analogously to individual TB-PE samples. As shown in **Figure S1**, no significant differences were observed in the average response or variability of a biological parameter (lactate release from M1 macrophages) between three pools (each comprising four distinct TB-PE samples) and four independent individual TB-PE samples (unpaired t-test, p = 0.115). These findings justify giving equal statistical weight to pools and individual samples in the lipidomic analysis (**Supplementary Tables I-II**).

### Local Synthesis of RvD5 by Monocytes and Expression of its Receptor GPR32 on Macrophages in Tuberculous Pleuritis

Our next objective was to investigate the potential origins of SPMs in the TB-PE. To address this, we analyzed the expression of key enzymes involved in SPM biosynthesis, namely arachidonic acid 5-lipoxygenase (5-LO, ALOX5), along with arachidonic acid 12- and 15-lipoxygenating paralogues (ALOX12, ALOX15, and ALOX15B)^22^, in peripheral blood mononuclear cells (PBMC) and pleural fluid mononuclear cells (PFMC) from TB patients. This analysis utilized publicly available single-cell RNA-Seq datasets (accession numbers HRA000910 and HRA00036). According to the literature^23^, lymphocytes (mainly T cells) dominate leukocyte infiltrates, while monocytic populations are underrepresented in TB-PE compared to blood (**Figure 2A)**. While monocytes from both compartments showed minimal expression of ALOX12, ALOX15, and ALOX15B, that of ALOX5 was significantly upregulated in pleural CD14⁺ cells compared to circulating monocytes (**Figure 2B, S2A**). In CD14⁺ monocytes specifically, ALOX5 expression increased ∼4.4-fold in PFMC versus PBMC (predicted means: ∼0.33 to ∼1.48), with no significant changes observed in other cell types (**Figure 2C, S2B**). This selective upregulation suggests that, within the context of pleural TB, pleural CD14⁺ cells are the primary mononuclear cell population responsible for local specialized pro-resolving mediators.

**Figure 2.**
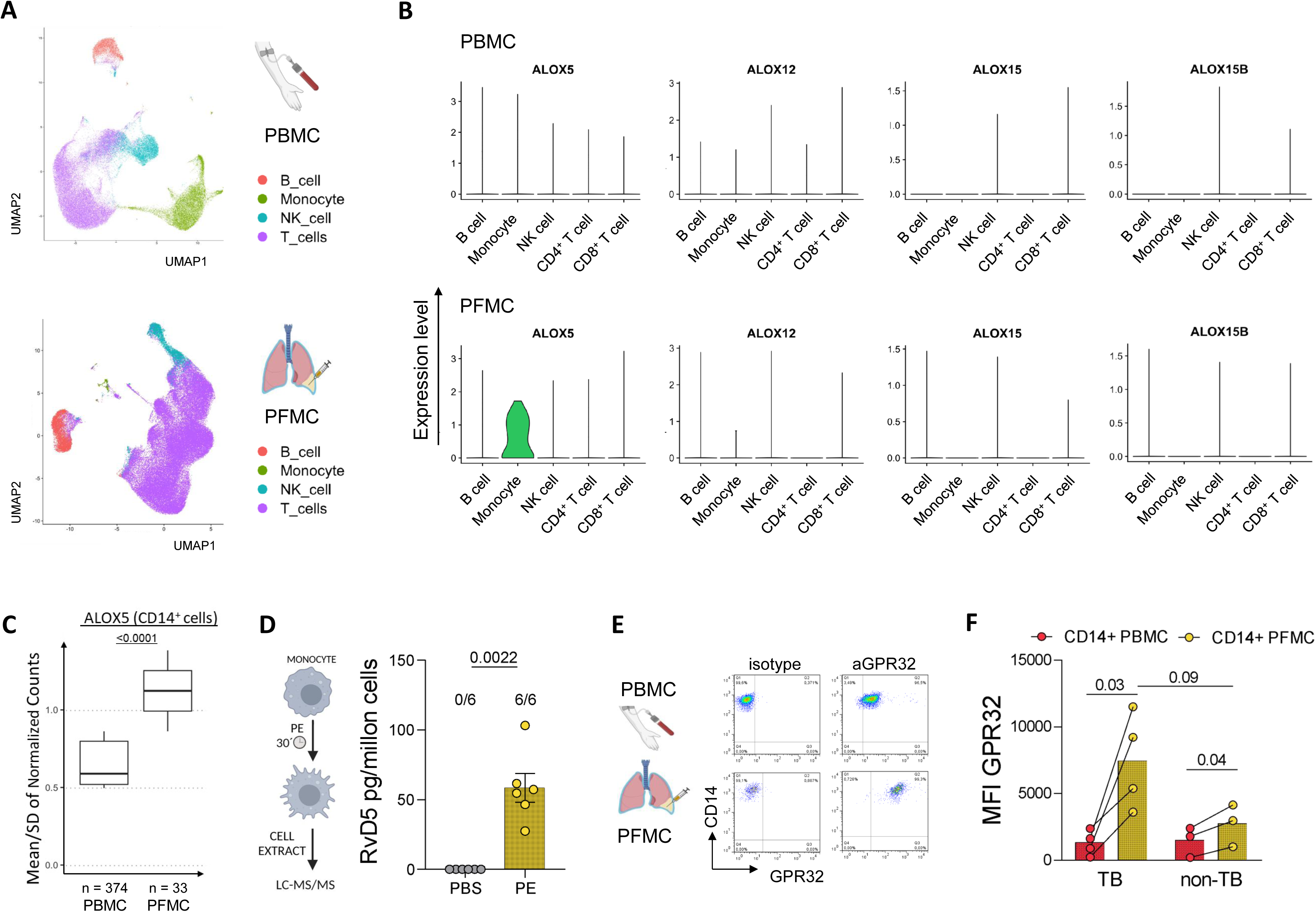
CD14^+^ cells synthesize RvD5 and its receptor GPR32 in tuberculous pleural effusions. **(A)** Single cell RNA-seq data visualized through a 2D UMAP plot: visualization of mononuclear cells from blood and pleural fluid of TB patients, with cells color-coded by major cell types (PBMC, n=5; PFMC, n=5). **(B)** Expression of enzymes involved in specialized pro-resolving mediator (SPM) biosynthesis. Violin plots depict the expression levels of key lipoxygenase genes (ALOX5, ALOX12, ALOX15, ALOX15B) across B cell, monocyte (CD14^+^), NK cell, and CD4 and CD8 T cell lineages. Data are shown for peripheral blood mononuclear cells (PBMCs; upper panels, n=5) and pleural fluid mononuclear cells (PFMCs; lower panels, n=6) from patients with TB. **(C)** Box-and-whisker plots represent the mean-to-variability ratio (μ/σ) of normalized ALOX5 expression in monocyte cells (CD14^+^) from TB patients. The ratio was calculated separately for PBMC (n=5 patients) and PFMC (n=4 patients). The total number of cells in each group is indicated on the x-axis. Each box summarizes the distribution of μ/σ values across patients within each group, with boxes showing the interquartile range (IQR), whiskers extending to 1.5×IQR, and center lines indicating median values. Statistical comparison was performed using a nested t-test. **(D)** Quantification of RvD5 in cellular extracts from healthy blood monocytes treated in vitro with either PBS 1x or TB-PE (n=6). The proportion of RvD5-producing monocytes out of a total of 6 independent donors is shown above each bar. Statistical analysis was performed using Fisher’s exact test. **(E)** Flow cytometry analysis was performed to compare GPR32 expression between CD14+ cells from peripheral blood and pleural effusions. Representative dot plots showing GPR32 expression on CD14^+^ cells from paired blood and TB pleural mononuclear cells. Representative flow cytometry dot plots show GPR32 expression on paired peripheral blood and pleural fluid CD14⁺ mononuclear cells from a TB patient. **(F)** GPR32 levels, expressed as median fluorescence intensity normalized to fluorescence-minus-one controls (MFI/FMO), on CD14⁺ cells from paired blood and pleural fluid samples. Data includes TB patients (n=4) and non-TB patients (n=3). Individual data points are shown as scatter plots; horizontal bars represent the mean ± SEM. Statistical comparisons were performed using a paired t-test (paired blood vs. pleural expression) and an unpaired t-test (TB vs. non-TB groups). PBMC: peripheral blood mononuclear cells; PFMC: pleural fluid mononuclear cells.

We also examined RvD5 levels in monocytes from blood of healthy donors exposed to the acellular fraction of TB-PE, compared with unexposed monocytes, to replicate the pleural environment encountered by newly recruited monocytes. After exposure, the monocytes were extensively washed, and intracellular RvD5 levels were measured. RvD5 was significantly detected in extracts from monocytes treated with the acellular fraction of TB-PE (**Figure 2D**). Together, these findings indicate that monocytes produce RvD5 in response to environmental factors in the pleural cavity of patients with pleural TB.

Building on these results, we next explore the potential involvement of the RvD5-GPR32 axis in pleural TB pathogenesis. Previous studies have shown that GPR32 is highly expressed on macrophages and its activation induces macrophage polarization from a pro-inflammatory to a pro-resolutive phenotype^24^. We first confirmed the presence of the RvD5 receptor, GPR32, on cells within the pleural cavity. Expression was significantly higher in pleural CD14^+^ cells than in their circulating counterparts, a pattern consistent in both TB and non-TB patients. Furthermore, GPR32 expression in pleural CD14^+^ cells was clearly increased in TB patients compared with non-TB patients **(Figure 2E-F)**. This demonstrates the receptor’s expression in cells locally exposed to RvD5.

### The RvD5-GPR32 axis mediates the inhibition of macrophage glycolysis by tuberculous pleural effusions

To determine whether RvD5 was responsible for the inhibition of macrophage glycolysis triggered by TB-PE, we conducted functional assays using commercially available RvD5. As previously shown^25^, compared to non-activated macrophages (M0), M1 macrophages typically exhibited high glycolytic activity by increasing lactate release (**Figure 3A**). To determine the contribution of RvD5 to the observed effect, we treated M1 macrophages with 17 and 33 nM RvD5, concentrations chosen to mirror those found in 20% TB-PE. Under these conditions, which reflect the physiological stoichiometry of the effusion, RvD5 alone significantly reduced lactate production, matching the inhibition achieved by TB-PE 20% v/v **(Figure 3A and S3A).** Notably, RvD5 treatment did not affect cell viability (**Figure S3B**). Given the pivotal role of HIF1α in the TB-PE-induced impairment of M1 macrophage functions^15^, we explored whether RvD5 influenced the upregulation of HIF1α in M1 cells. Our findings revealed that RvD5 reduced HIF1α expression in M1 macrophages (**Figure 3B**). Similarly, RvD5 suppressed lactate release when M1 macrophages (activated by IFNγ) were challenged with irradiated *M. tuberculosis* (**Figure 3C**).

**Figure 3.**
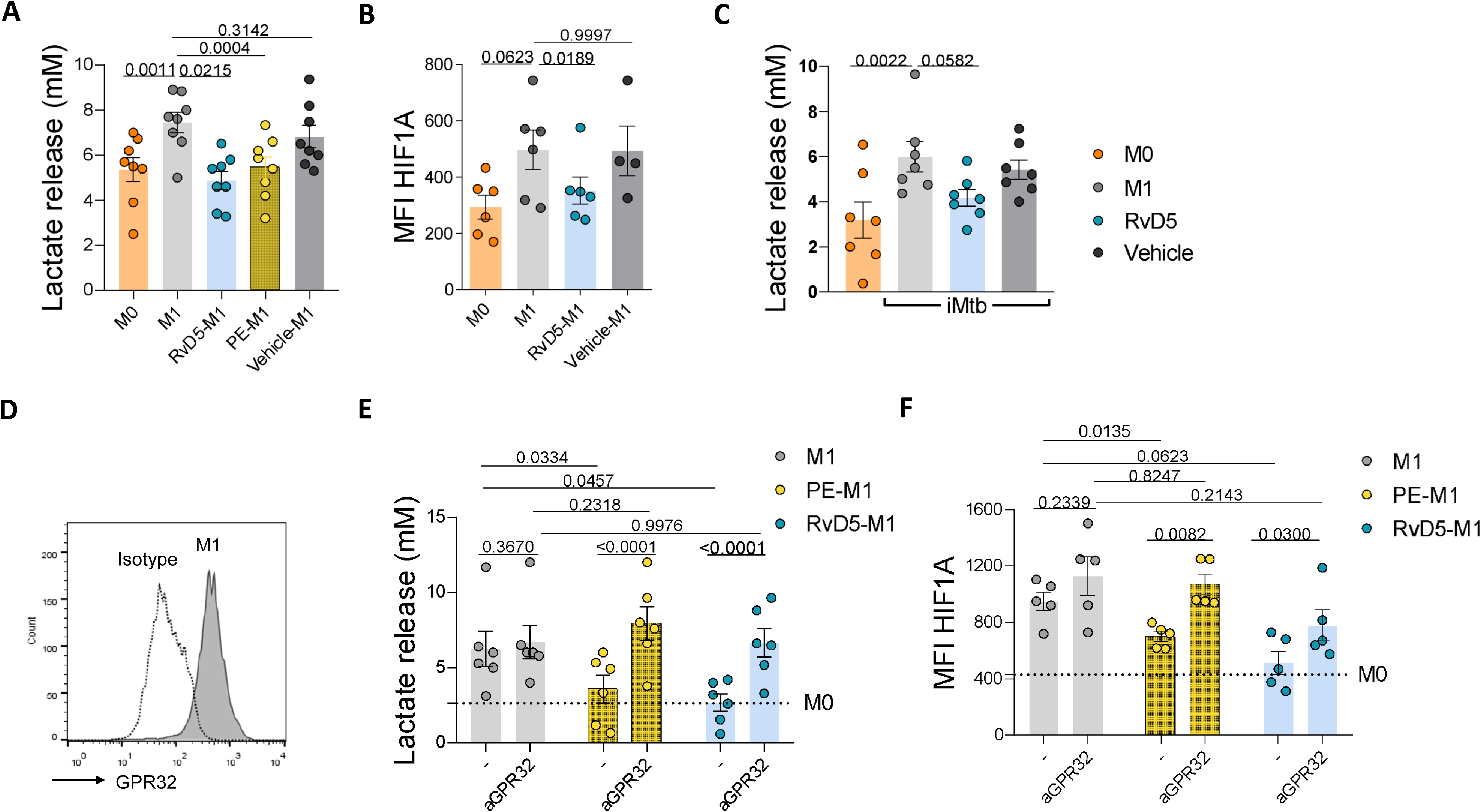
The RvD5-GPR32 axis mediates the inhibition of macrophage glycolysis induced by TB pleural effusions. **(A)** Lactate release from unstimulated (M0) or LPS/IFNγ-activated (M1) macrophages treated for 24h with RvD5 (33 nM), 20% v/v TB pleural effusion (TB-PE), or vehicle (ethanol) (n=8). Data were analyzed by one-way ANOVA with Dunnett’s multiple comparisons test. **(B)** HIF1α expression in M0 or M1 macrophages exposed to RvD5 (n=6) or vehicle (n=4). Data was analyzed using a mixed-effects model with Sidak‘s multiple comparisons test. **(C)** Lactate release from unstimulated (M0) or IFNγ-activated (M1) macrophages exposed to irradiated *M. tuberculosis* (iMtb) and treated with RvD5 or vehicle for 24h (n=7). Data was analyzed by one-way ANOVA with Dunnett’s multiple comparisons test. **(D)** Representative flow cytometry histogram showing GPR32 surface expression on LPS/IFNγ-stimulated (M1) human macrophages, with an isotype control. **(E)** Lactate release from LPS/IFNγ-stimulated M1 macrophages pre-treated with a neutralizing anti-GPR32 antibody (αGPR32) or isotype control before addition of RvD5 or 20% v/v TB-PE (n=6). **(F)** HIF1α expression in LPS/IFNγ-stimulated M1 macrophages pre-treated with αGPR32 or isotype control before addition of RvD5 or 20% v/v TB-PE (n=5). The horizontal dotted line indicates the mean value from untreated M0 macrophages. Data in (E) and (F) were analyzed by two-way ANOVA. For the factor of antibody treatment, Holm-Sidak’s multiple comparisons test was applied (αGPR32 vs. isotype). For the stimulation condition factor, Tukey’s multiple comparisons test was applied (M1 vs. PE-M1 vs. RvD5-M1). All quantitative data are presented as scatter plots showing individual replicates with mean ± SEM.

Given the high expression of GPR32 observed in CD14^+^ pleural cells, we investigated whether the inhibitory effects of exogenous RvD5 are mediated through this receptor. First, in line with the literature^26^, we confirmed GPR32 expression in M1 macrophages (**Figure 3D**). Next, we observed that blocking GPR32 with neutralizing anti-GPR32 antibodies fully restored lactate release in M1 macrophages exposed to either RvD5 or TB-PE (**Figure 3E**), indicating that GPR32 signaling by TB-PE lipids was sufficient to alter the metabolism of M1 macrophages. Moreover, blockade of GPR32 also restored HIF1α levels in both RvD5 and PE-treated M1 macrophages (**Figure 3F)**.

These findings collectively demonstrate that RvD5 inhibits glycolysis in M1 macrophages through a GPR32-dependent mechanism, suppressing HIF1α activity.

### Activation of the RvD5-GPR32 axis reduces glycolysis without affecting mitochondrial bioenergetics in M1 macrophages

To investigate if the TB-associated microenvironment affects the metabolic profile of pleural monocyte/macrophages, we compared the *ex vivo* metabolic profile of CD14^+^ cells from TB-PE with that of paired circulating monocytes. This was achieved by monitoring changes in protein translation upon treatment with oligomycin and/or 2-deoxy-D-glucose using the SCENITH assay^27^. Our findings indicated a reduced glycolytic capacity in cells residing in the TB pleural cavity compared to blood cells (**Figure 4A**). Interestingly, pleural macrophages from TB patients undergoing antibiotic treatment exhibited a significant increase in glycolysis compared to those from untreated patients (**Figure 4A**). Our previous work established that circulating monocytes from TB patients have higher glycolytic metabolism than those from healthy individuals^28^. Here, we extended these observations to circulating monocytes from pleural TB patients, in whom glycolytic activity showed a decreasing trend upon antibiotic treatment **(Figure 4A**).

**Figure 4.**
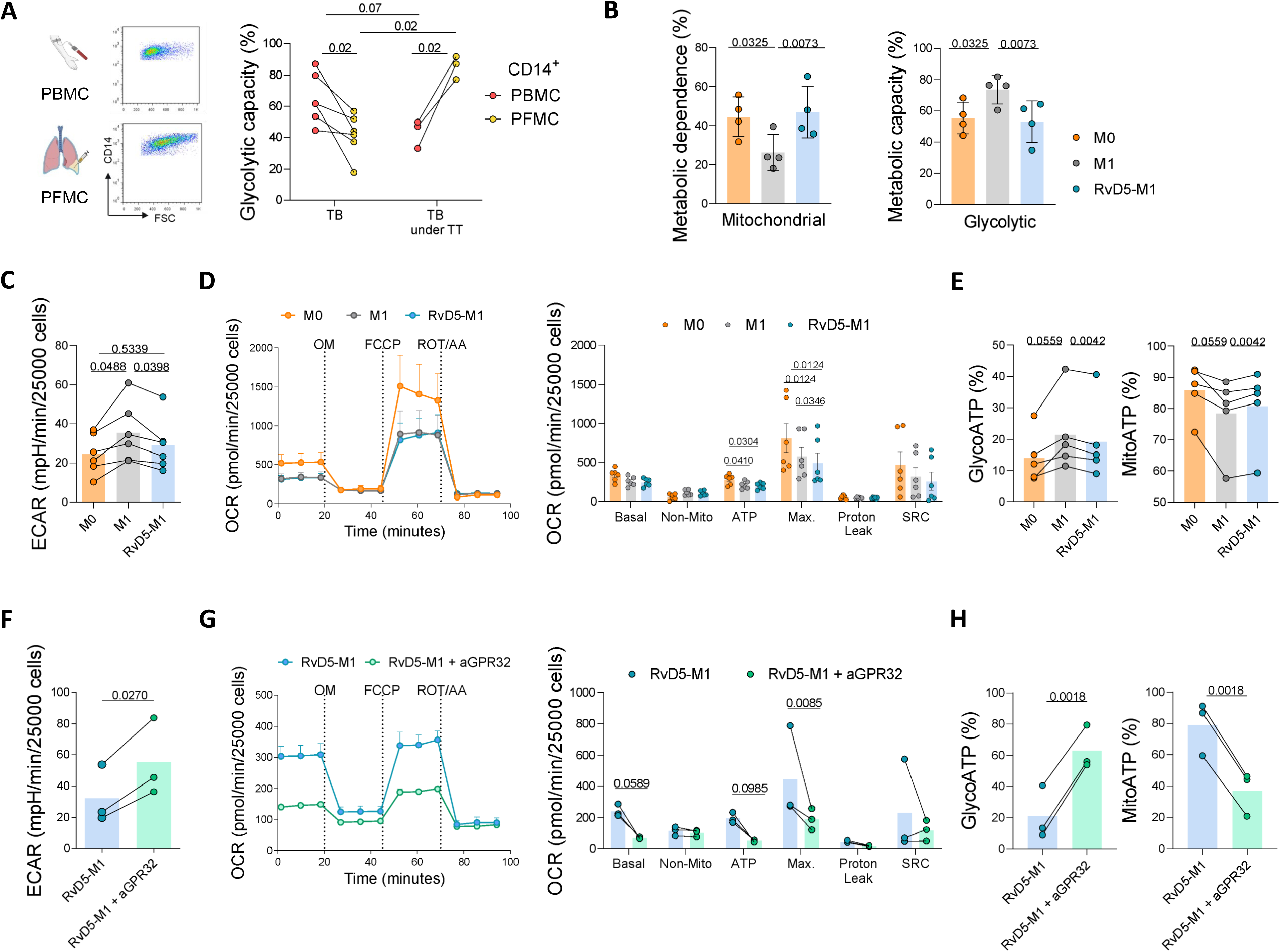
Activation of the RvD5-GPR32 Axis Reduces Glycolysis Without Affecting Mitochondrial Bioenergetics in M1 macrophages. **(A)** Basal glycolysis measured via SCENITH in CD14⁺ cells from paired blood mononuclear cells (PBMC) and pleural fluid mononuclear cells (PFMC) of untreated TB patients (n=5) and treated TB patients (n=3). Statistical analysis: paired t-test for PFMC vs. PBMC within each group; Mann–Whitney test for untreated vs. treated TB patients. **(B)** Mitochondrial dependence and glycolytic capacity analyzed via SCENITH in M0, M1, and RvD5-treated M1 macrophages (n=4). Data was analyzed using one-way ANOVA with Dunnett’s multiple comparisons test. **(C)** Extracellular acidification rate (ECAR) measured in M0 and M1 macrophages treated or not with RvD5 (n=6). Data was analyzed using one-way ANOVA with Sidak’s multiple comparisons test. **(D)** Oxygen consumption rate (OCR) traces and respiration parameters for M0 and M1 macrophages treated or not with RvD5. Respiratory parameters include basal respiration, ATP-linked respiration, proton leak, maximal respiration, spare respiratory capacity (SRC), and non-mitochondrial respiration. Representative trace from one of six independent experiments is shown; bar graphs show means ± SEM (n=6). Data was analyzed using two-way ANOVA with the Benjamini, Krieger and Yekutieli two-stage linear step-up procedure. **(E)** Percent contributions of ATP from mitochondrial oxidative phosphorylation (MitoATP) and glycolysis (GlycoATP) in M0 and M1 macrophages treated or not with RvD5 (n=5). Data was analyzed using one-way ANOVA with Sidak’s multiple comparisons test. **(F)** ECAR in M1 macrophages treated with RvD5, with or without neutralizing anti-GPR32 antibody (αGPR32) (n=3). Statistical analysis: paired t-test. **(G)** OCR traces and respiration parameters for RvD5-treated M1 macrophages in the presence or absence of αGPR32 (n=3). Data were analyzed using two-way ANOVA with Benjamini, Krieger, and Yekutieli procedure. **(H)** Percent contributions of MitoATP and GlycoATP in RvD5-treated M1 macrophages in the presence or absence of αGPR32 (n=3). Statistical analysis: paired t-test. All quantitative data are presented as scatter plots showing individual donors; means ± SEM are indicated. Data obtained from Seahorse experiments were normalized based on the area covered by the cells, and a scale factor of 25,000 was used.

Additionally, we evaluated the impact of RvD5 on the relative contributions of metabolic pathways to the cellular energy metabolism of M1 macrophages using SCENITH. According to the expected glycolytic profile of M1 activation program^29^, M1 macrophages exhibited a lower reliance on OXPHOS alongside an increase in the glycolytic capacity compared to unstimulated cells (M0) (**Figure 4B**). However, this metabolic shift in M1 macrophages was impaired upon addition of exogenous RvD5 **(Figure 4B and Figure S3 and S4)**. Of note, while TB-PE has been shown to promote lipid body accumulation in macrophages^13,14^, RvD5 did not induce lipid body formation, at least at the tested doses **(Figure S5**). These findings indicate that RvD5 reduces the glycolytic activity of M1 macrophages independently of lipid body accumulation, unlike TB-PE.

To examine the effect of RvD5 on the OXPHOS metabolic state in macrophages, we performed the Seahorse XF Cell Mito Stress Test in M0, M1, and RvD5-treated M1 cells. To characterize the overall energy phenotypes of these macrophages, basal oxygen consumption rate (OCR) was plotted against extracellular acidification rate (ECAR). The resulting metabolic phenogram revealed that RvD5 shifts the energy metabolism of M1 macrophages towards a less glycolytic profile without impacting aerobic metabolism (**Figure S6A**). This shift is further reinforced by a small but significant decrease in basal ECAR levels observed in RvD5-treated M1 cells compared to untreated M1 cells (**Figure 4C**), aligning with the metabolic characterization from SCENITH and lactate measurements described earlier (**Figure 3A and 4B**).

Analysis of respiratory parameters indicates substantial reductions in ATP-linked OCR and maximal respiratory capacity in M1 macrophages compared to M0 cells. While RvD5 treatment did not reverse these M1-induced deficits, it caused a further, specific reduction in maximal respiratory capacity (**Figure 4D**).

To further evaluate intracellular ATP production, we measured the basal ECAR and OCR to calculate ATP production rates from glycolysis (GlycoATP) and mitochondrial OXPHOS (MitoATP). M1 macrophages exhibited higher ATP production rates from glycolysis compared to RvD5-treated M1 macrophages (**Figure S6A, C-D**), with no differences in ATP production from mitochondrial respiration (**Figure S6A-B and D**). Consistent with SCENITH findings, the relative contribution of MitoATP to total ATP production increased, while the contribution of GlycoATP decreased in RvD5-treated M1 macrophages compared to untreated M1 cells (**Figure 4E**). Furthermore, in line with these results, M1 macrophages showed lower mitochondrial mass than control cells, but this parameter remained unchanged upon RvD5 treatment **(Figure S7A-B)**. Similarly, transmission electron microscopy (TEM) revealed no differences in mitochondrial morphology between RvD5-treated and untreated M1 macrophages (**Figure S7C**). These findings suggest that RvD5 reduces glycolysis without increasing aerobic metabolism, thereby promoting a less energetically active metabolic state.

Finally, we assessed whether specific inhibition of GPR32 could restore M1 metabolism in RvD5-treated macrophages. Our results revealed that blocking GPR32 led to a notable shift in RvD5-M1 cells, driving them towards heightened glycolysis, as evidenced by an increase in basal ECAR (**Figure 4F and Figure S8A and C**). Notably, key respiratory parameters, including basal OCR and maximal respiratory capacity, were significantly diminished in RvD5-M1 macrophages following GPR32 inhibition (**Figure 4G**), suggesting a compensatory reduction in OXPHOS to accommodate the increased glycolysis. These findings were further corroborated by ATP production analysis, which yielded a significant increase in GlycoATP levels and a corresponding decrease in MitoATP levels upon GPR32 inhibition (**Figure S8B-D**). Additionally, the relative contribution of GlycoATP to total ATP production increased, while that of MitoATP decreased, compared with cells treated with the isotype control (**Figure 4H**). This metabolic reprogramming highlights a pronounced shift towards glycolytic reliance at the expense of the OXPHOS activity.

Collectively, these results underscore the critical role of the RvD5-GPR32 axis in suppressing glycolytic activity in M1 macrophages, maintaining a hypometabolic state without increasing OXPHOS flux.

### RvD5 boosts a pro-resolving phenotype in M1 macrophages at the expense of its mycobacterial control potential

Since RvD5 is an SPM, we assessed the acquisition of pro-resolutive features by M1 macrophages exposed to RvD5. M1 macrophages displayed low efferocytic activity, which was significantly enhanced by RvD5 treatment, as evidenced by an increased engulfment of apoptotic neutrophils (**Figure S9A**). This improvement was accompanied by an upregulation of MerTK, a major efferocytic receptor^30^, following RvD5 treatment (**Figure S9B**), along with an increased release of the anti-inflammatory cytokine IL-10 (**Figure S9C**). In contrast, the expression levels of other efferocytosis receptors, such as CD14^31^ and the scavenger receptor CD36^32^, remained unchanged between M1 macrophages and RvD5-treated M1 cells (**Figure S9D**).

Then, we examined the capacity of M1 macrophages to engulf viable Mtb. There was no significant change in the initial bacterial uptake between RvD5-treated M1 macrophages and untreated cells (**Figure 5A**). However, RvD5 treatment lowers IL-1β secretion by M1 macrophages in response to Mtb infection and impaired strongly their microbicidal activity (**Figure 5B-C)**. Additionally, while Mtb-infected macrophages displayed a higher LC3-II/LC3-I ratio compared with uninfected cells, this induction was reduced when M1 cells were exposed to RvD5 (**Figure 5D**). Consistent with this observation, punctate LC3 labeling was predominantly seen in Mtb-infected macrophages relative to RvD5-treated, Mtb-infected macrophages (**Figure 5E)**. Together, these results demonstrate that RvD5 prevents the induction of autophagy in Mtb-infected macrophages, thereby impairing a key mechanism through which macrophages control bacterial growth^33^.

**Figure 5.**
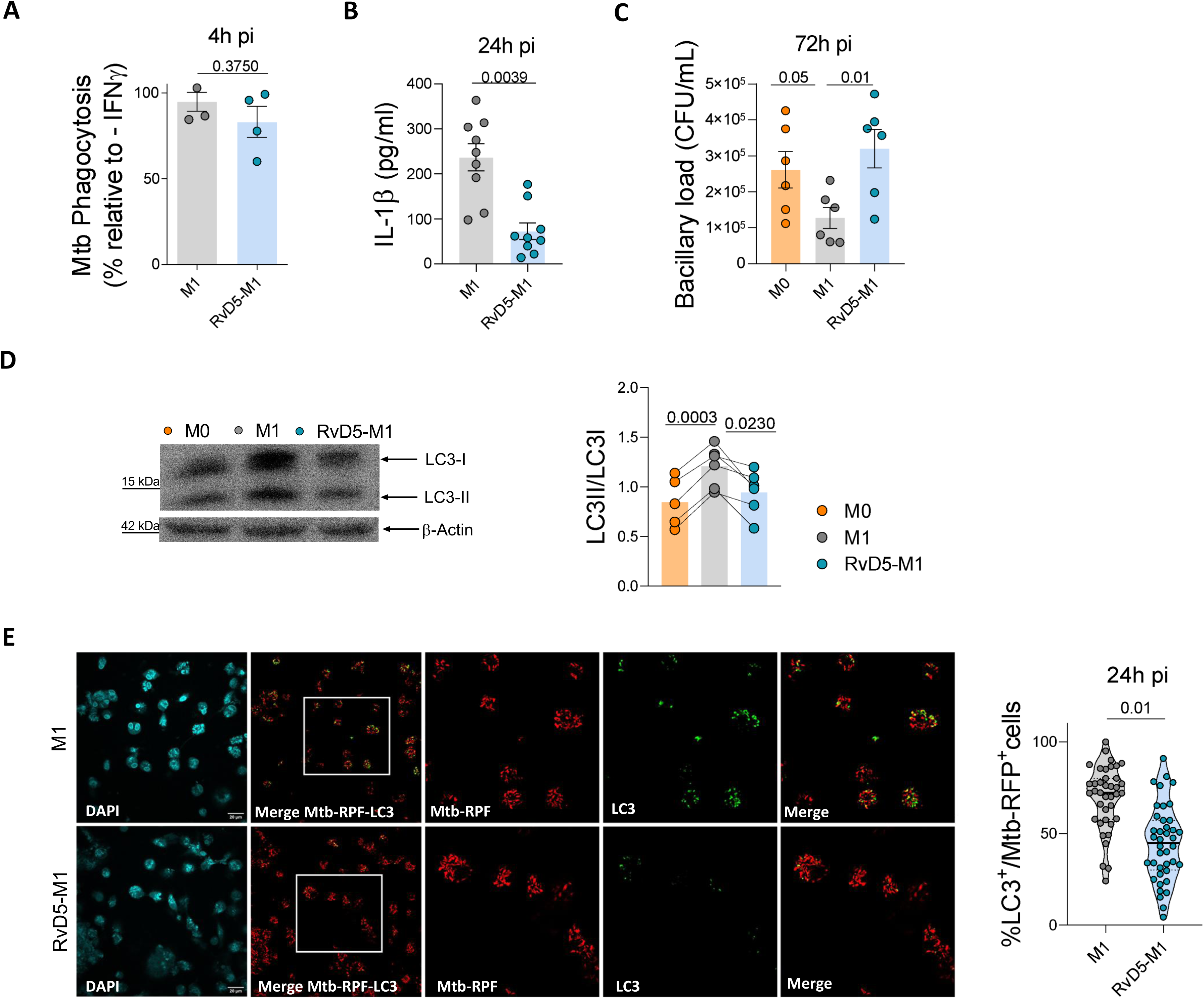
RvD5 Impairs Mycobacterial control by M1 Macrophages. **(A)** Initial uptake of *Mtb*. Unstimulated (M0) or IFNγ-activated (M1) macrophages, treated or not with RvD5, were infected with *Mtb*-RFP. The percentage of RFP^+^ cells was quantified 4 h post-infection. Results are expressed relative to uptake by unstimulated cells (n=4). Statistical analysis: Wilcoxon matched-pairs signed-rank test comparing M1 macrophages with vs. without RvD5. **(B)** IL-1β levels. Supernatants collected 24 h post-infection were analyzed by ELISA (n=9). Statistical analysis: Wilcoxon matched-pairs signed-rank test comparing M1 macrophages with vs. without RvD5. **(C)** Mycobacterial growth. Colony-forming units (CFU) were quantified 72 h post-infection in unstimulated or M1 macrophages with and without RvD5 (n=6). Data was analyzed by one-way ANOVA followed by Holm-Sidak’s multiple comparisons test. All quantitative data are presented as scatter plots of individual donors, with means ± SEM shown. **(D)** M0 macrophages remained uninfected and unpolarized. M1 macrophages were pretreated with RvD5 or not prior to infection with Mtb for 6 h. Cell lysates were subjected to western blotting to detect LC3-I, LC3-II, and β-actin as a loading control. The Right panel shows the densitometric quantification of the LC3-II/LC3-I ratio; individual data points from the same donor are connected by lines to reflect the paired experimental design (n=6 donors per group). Data were analyzed by one-way ANOVA with Tukey’s multiple comparisons test. **(E)** Representative confocal micrographs show M1 macrophages infected with *Mtb*-RFP for 24 h in the presence or absence of RvD5. Nuclei are stained with DAPI (blue), and the autophagy marker LC3 is stained with an Alexa Fluor 488-conjugated antibody (green). The percentage of infected cells (RFP^+^) exhibiting LC3 puncta per field was quantified. Statistical analysis: Nested t-test from four independent experiments, analyzing 10 images per condition.

Consistent with the role of autophagy-dependent mycobacterial degradation in enhancing antigen presentation to T cells^34,35^, we observed reduced CD4 T cell activation in antigen-presenting cell (APC)-dependent responses against irradiated *Mtb* (iMtb) when RvD5 was added to the cocultures (**Figure S10A–D**). Notably, this effect was absent in APC-independent T cell activation assays (**Figure S10E–G**), providing additional evidence that RvD5 specifically compromises macrophage function. Interestingly, rapamycin treatment reversed the RvD5-mediated inhibition of T cell activation, indicating that pharmacological restoration of autophagy is sufficient to improve antigen presentation (**Figure S10H-I**). Together, these data directly link the effect of RvD5 on autophagy to impaired T cell responses.

Altogether, our results indicate that the omega-3-derived lipid mediator RvD5 reprograms the proinflammatory profile of M1 macrophages toward a more pro-resolving, immunomodulatory phenotype.

### HIF1α stabilization restores M1 macrophage features inhibited by the GPR32-RvD5 axis

To investigate whether the inhibition of HIF1α mediates the inhibition of the microbicidal activity of M1 macrophages induced by the RvD5-GPR32 axis, we stabilized HIF1α expression by using Dimethyloxalylglycine (DMOG). Remarkably, HIF1α stabilization in RvD5-treated M1 macrophages restored their impaired lactate and IL-1β release (**Figure 6A-B**), as well as their microbicidal activity, evidenced by reduced bacillary loads and a higher localization of Mtb in compartments marked by the late endosomal marker, LAMP2 (**Figure 6C-D)**. Conversely, DMOG treatment diminished the pro-resolution features induced by RvD5, including enhanced efferocytic capacity, upregulated MerTK expression, and increased IL-10 production (**Figure S11**).

**Figure 6.**
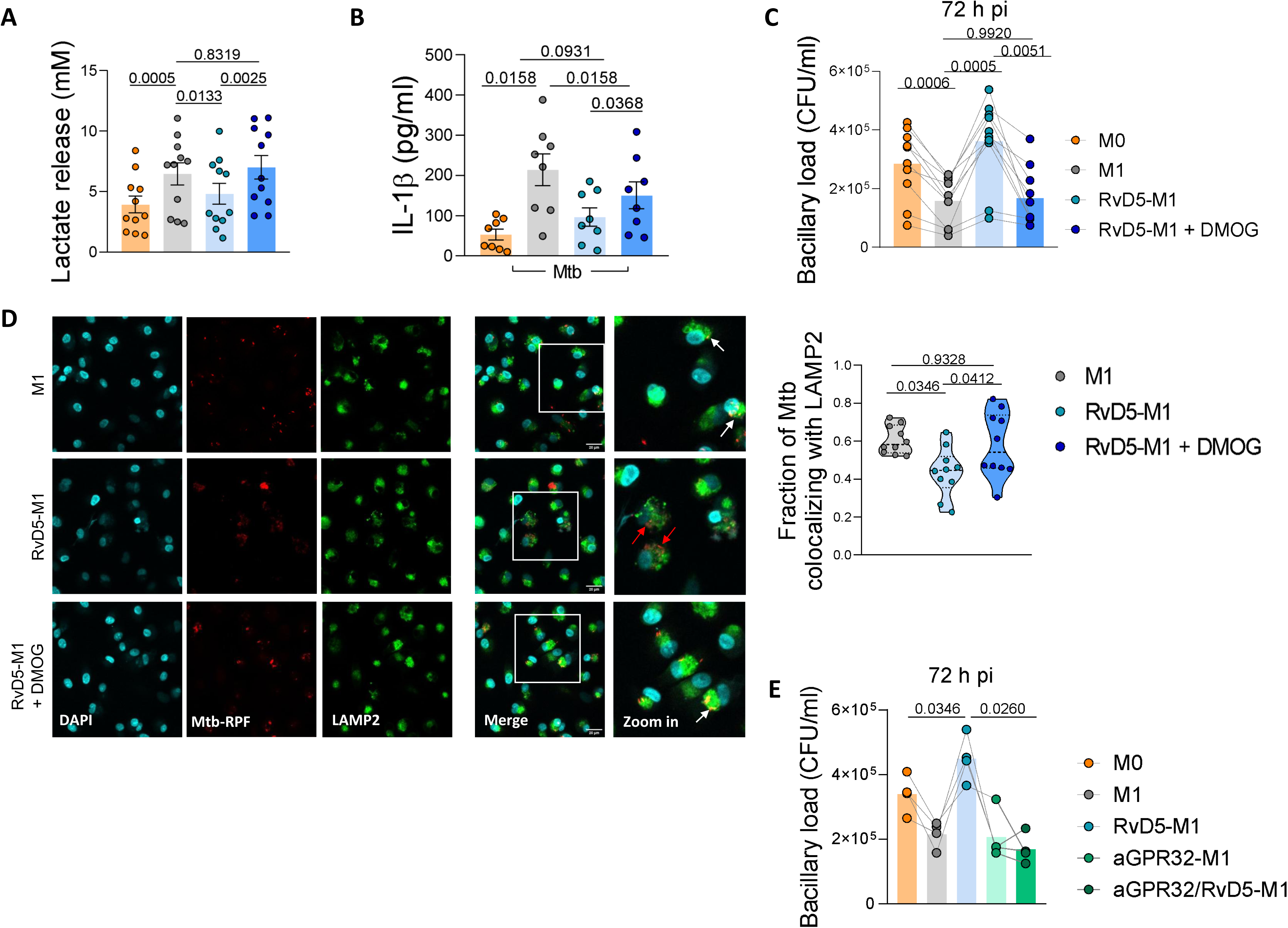
Chemical HIF1α Stabilization Restores RvD5-mediated M1 Macrophage metabolism and function. **(A)** Lactate release from unstimulated (M0) or LPS/IFNγ-activated (M1) macrophages treated with RvD5 in the presence or absence of the HIF-1α stabilizer DMOG for 24 h (n=11). **(B–D)** IFNγ-activated macrophages (M1), treated or not with RvD5 and/or DMOG, were infected with *Mtb*. **(B)** IL-1β levels measured by ELISA in supernatants collected 24 h post-infection (n=8). **(C)** Mycobacterial growth assessed by colony-forming units (CFU) quantified 72 h post-infection (n=10). For panels A–C, data were analyzed by one-way ANOVA followed by Sidak’s multiple comparisons test. All quantitative data are presented as scatter plots of individual donors, with means ± SEM. **(D)** Colocalization of *Mtb*-RFP with LAMP2. Representative confocal micrographs show macrophages infected with *Mtb*-RFP for 24 h. Nuclei are stained with DAPI (blue), and the late endosome marker LAMP2 is stained with an Alexa Fluor 633-conjugated antibody (cyan). The lower panels show magnified views of the boxed areas (scale bar = 10 μm). The right panel shows Manders colocalization coefficients for the fraction of *Mtb* signal overlapping with LAMP2 staining. Data was from a representative experiment (10 images per condition) and was analyzed by one-way ANOVA followed by Tukey’s multiple comparisons test. **(E)** Mycobacterial growth assessed by CFU quantified 72 h post-infection in M1 macrophages treated with RvD5, with or without neutralizing anti-GPR32 antibody (αGPR32, n=5).

Furthermore, blocking GPR32 in the presence of RvD5 significantly improved control of intracellular bacterial growth (**Figure 6E**), providing direct evidence that RvD5 signals through GPR32 to impair microbicidal activity.

Together, these findings demonstrate that the RvD5-GPR32-HIF1α axis suppresses macrophage microbicidal functions and promotes susceptibility to *Mtb*. Importantly, these effects can be reversed by pharmacological stabilization of HIF1α.

## Discussion

Pleural TB presents unique challenges for healthcare providers, particularly concerning its treatment. Effective host defense against TB involves infiltration of peripheral blood cells into the pleural space, leading to accumulation of immune cells and soluble factors, including lipid mediators, in tuberculous pleural fluid. SPMs play a critical role in limiting inflammation and preventing collateral damage to healthy tissues. However, it is essential to carefully regulate the potential adverse effects of SPMs on the microbicidal functions of inflammatory macrophages to achieve a balanced approach that supports adequate pathogen clearance while minimizing excessive inflammation. While previous studies have highlighted the beneficial roles of SPMs in enhancing the microbicidal activities of macrophages and neutrophils^21,36–38^, our study is the first to establish the involvement of the RvD5-GPR32-HIF1α axis in TB. We demonstrate that RvD5, present in pleural TB, activates GPR32, thereby inhibiting HIF1α-dependent microbicidal activity in macrophages.

Our study centers on two pivotal findings. First, we identified the RvD5-GPR32 axis as a key mediator of macrophage glycolysis suppression in TB pleural effusions. We demonstrated that RvD5 is not only present in pleural effusions but can also be produced by monocytes exposed to the acellular fraction of these effusions. Notably, CD14^+^ cells in the tuberculous pleural cavity constitute the primary mononuclear cell population responsible for local production of SPMs, such as resolvins. The accumulation of RvD5 within the pleural cavity correlates with reduced glycolytic capacity in CD14^+^ cells compared to their circulating counterparts. This suggests that upon entering the pleural cavity, phagocytes undergo a metabolic shift characterized by attenuated glycolytic activity, influenced by the local microenvironment. In line with this, in vitro exposure of macrophages to RvD5 led to a similar reduction in glycolysis. In this work, we used M1-polarized cells to model the *in vitro* effects of pleural TB, given that pleural effusions from patients with active TB contain high levels of IFN-γ– in fact, IFN γ has been proposed as a diagnostic marker for tuberculous pleural effusions compared to other pleural diseases^39,40^. We acknowledge that this represents a simplified view of the complex, mixed macrophage phenotypes present in the pleural cavity. However, key findings (such as GPR32 expression and metabolic changes) were validated directly in CD14⁺ cells isolated from pleural effusions, irrespective of their polarization status. This independent validation strengthens the physiological relevance of our observations.

We show that pleural macrophages from TB patients undergoing antibiotic treatment exhibit a significant increase in glycolytic activity compared to those from untreated patients. This observation indicates the presence of active mechanisms within the pleural cavity that suppress long-term protective immune responses, likely modulated by SPMs. Such findings offer a potential explanation for the frequent relapses observed in untreated pleural TB patients, suggesting that while glycolysis in macrophages is locally suppressed during active disease, systemically, monocytes in TB patients maintain a dysregulated glycolytic profile, as we recently reported^28^. This dichotomy supports the notion of contrasting metabolic environments at local and systemic levels. Even though using drugs that alter metabolism in all cells might have deleterious implications for patients, identifying strategies to do so selectively in macrophages has important clinical implications. We propose that upon entering the pleural cavity, monocytes upregulate ALOX5 transcript levels, thereby promoting local biosynthesis of SPMs. These SPMs, in turn, may modulate the metabolism of inflammatory macrophages, shifting them toward reduced glycolysis. This immunosuppressive metabolic state, however, appears reversible with TB treatment, underscoring the dynamic interplay between local immune modulation and systemic immune dysregulation in pleural TB.

Several studies have identified associations between ALOX5 polymorphisms and increased susceptibility to TB^41,42^ or the development of multisystem TB^43^. These studies primarily focus on the role of ALOX5 in synthesizing pro-inflammatory lipid mediators, such as leukotrienes. In contrast, our findings identify RvD5 as a novel and underappreciated player in this context. Supporting our observations, ALOX5-deficient mice have been shown to exhibit increased resistance to Mtb infection^44^. Interestingly, ALOX5 expression has been implicated in both pro-inflammatory and immunosuppressive outcomes across various pathological contexts. For example, in rheumatoid arthritis, increased ALOX5 expression has been associated with harmful inflammation by driving pyroptosis in CD4^+^ T cells^45^. Conversely, in gliomas, ALOX5 promotes an immunosuppressive phenotype by facilitating M2 polarization in macrophages^46^. These diverse outcomes are likely the eclectic array of products ALOX5 can generate, depending on substrate availability. Indeed, ALOX5 can metabolize arachidonic acid, EPA, or DHA to produce oxylipins with distinct biological functions, including 5-HETE, lipoxins, and resolvins^47^. This substrate-driven variability highlights the complexity of ALOX5’s involvement in TB and other diseases. It suggests that its effects may change depending on microenvironmental factors, such as lipid availability, cellular metabolic states, and the balance between pro-inflammatory and pro-resolving mediators. Understanding these dynamics is critical for fully elucidating the dual roles of ALOX5 and for considering therapeutic strategies that target its metabolic pathways in TB and beyond, and inhibition of ALOX5 might have undesired effects due to its pleiotropic function. On the positive side, GPR32 belongs to the most intensively studied drug target family, mainly because of its substantial involvement in human pathophysiology and its pharmacological tractability^48^.

Controlling excessive inflammation in the pleural space is crucial to prevent tissue damage and maintain host-pathogen balance. While RvD5 has been shown to impair the microbicidal functions of macrophages, other SPMs prevalent in pleural TB, such as PDx and MaR1, do not exhibit these detrimental effects^21^. This underscores an opportunity to selectively enhance SPMs that preserve microbicidal activity while minimizing tissue damage, thereby avoiding lipid mediators such as RvD5. For instance, in tuberculous meningitis, elevated concentrations of SPMs in cerebrospinal fluid have been associated with improved patient survival^49^. These results support the idea of targeted interventions to harness the benefits of SPMs while mitigating the risks.

The literature extensively explores the balance between “proinflammatory” and “protective” lipid mediators, often manipulated through dietary supplementation with omega-3 fatty acids (n-3 PUFAs, precursors of protective lipids) versus omega-6 fatty acids (n-6 PUFAs, mostly precursors of proinflammatory lipids). However, many studies lack a detailed examination of the mechanisms of action for individual lipids. The outcomes of such supplementation studies have been inconsistent, ranging from detrimental effects, such as the impairment of murine macrophage responses to Mtb with omega-3 supplementation^50,51^, to beneficial outcomes where n-3 PUFAs enhance immune responses against Mtb^50,52^. These conflicting findings underscore the need for further research to delineate the specific roles and mechanisms of individual lipid mediators in the context of TB.

Although some reports propose that certain SPMs, such as RvD1^53^ or MaR1^54^, can enhance cellular respiration in neutrophils and macrophages, our findings did not demonstrate that RvD5 increases absolute mitochondrial respiration in macrophages stimulated with LPS and IFNγ. This discrepancy may be due to the strong metabolic imprinting induced by LPS and IFNγ, which results in decreased OXPHOS that RvD5 cannot counteract. Alternatively, the effect of RvD5 may be limited to reducing glycolysis without activating compensatory energetic mechanisms. These insights further support the complex, context-dependent nature of SPM-mediated metabolic regulation, warranting more in-depth studies to fully understand their roles and therapeutic potential in TB.

Our second key finding reveals that activation of GPR32 by RvD5 inhibits HIF1α-dependent microbicidal activity in macrophages, shedding light on a critical pathway that modulates immune responses in TB. Previously, we demonstrated that lipids extracted from the acellular fraction of TB pleural effusions suppress HIF1α expression, leading to reduced glycolytic and microbicidal phenotypes in macrophages^15^. The present study goes beyond our previous results, as we now show that RvD5 is explicitly a central lipid mediator driving this effect through GPR32 binding. While GPR32 is the best-characterized receptor for RvD5, emerging evidence suggests that RvD5 may also interact with GPR101, which is also expressed in macrophages^53^. As GPR101 is constitutively active, any role for RvD5 would likely involve enhancing or fine-tuning its basal signaling rather than classical activation. The potential contribution of this alternative receptor to the observed metabolic effects was not explored in this study and remains an open avenue for future research.

Our results demonstrate that RvD5 impairs Mtb clearance, at least partially, by limiting the induction of autophagy in infected macrophages. Importantly, this suppression of autophagy could be reversed by stabilizing HIF1α, consistent with previous findings that HIF1α promotes autophagy in cancer cells^54^.

RvD5 also diminished IL-1β levels in Mtb-infected macrophages. IL-1β is essential for host control of Mtb: genetic mouse models lacking IL-1 signaling exhibit high mortality and increased bacterial burdens^55^, and clinical blockade of IL-1 with anakinra increases TB reactivation risk^56^. Critically for our study, *Il1b* transcription is regulated by HIF-1α-dependent glycolytic reprogramming upon Mtb infection^57,58^, and IL-1β itself induces autophagy leading to mycobacterial killing^57^. Thus, the RvD5-mediated reduction of both HIF-1α and IL-1β may cooperatively impair autophagy and bacterial clearance. Of note, it has been previously shown that *Mtb* can inhibit IFNγ-induced autophagy via an ALOX5-dependent pathway involving Mir155-Mir31-WNT-SHH signaling^59^. However, the role of RvD5 was not investigated in that study. Interestingly, unlike RvD5, other resolvins such as RvD1 promote autophagy in various contexts^60,61^. These findings emphasize the functional specificity of different SPMs and caution against generalizing biological functions across the resolvin family.

In conclusion, our study provides valuable insights into the pathophysiology of pleural TB by uncovering RvD5 as a critical lipid mediator that modulates macrophage metabolism and function through GPR32. This work underlines the importance of understanding the specific mechanisms by which lipid mediators influence immune cell metabolism and TB severity. These results pave the way for the development of targeted therapeutic strategies that aim to balance immune regulation, preserve microbicidal activity, and mitigate the detrimental effects of excessive inflammation in TB.

## Material and methods

### Human Subjects

Buffy coats from healthy donors were prepared at the Centro Regional de Hemoterapia Garrahan (Buenos Aires, Argentina) according to the institutional guidelines (Resolution No. CEIANM-664/07). Informed consent was obtained from each donor before blood collection.

Pleural effusions (PE) were obtained by therapeutic thoracentesis by physicians at the Hospital F. J Muñiz (Buenos Aires, Argentina). The diagnosis of TB pleurisy was based on a positive Ziehl–Neelsen stain or Löwenstein–Jensen culture from PE and/or histopathology of pleural biopsy, and was further confirmed by an Mtb-induced IFNγ response and an adenosine deaminase-positive test^10^. None of the patients had multidrug-resistant TB. For this study, we used acellular fractions of PE stored at-80°C for lipid analysis, and fresh paired pleural fluids and blood mononuclear cells (PFMC and PBMC) for assessment of metabolic profiles and GPR32 expression. PE samples were divided by etiology. For the lipidomics analysis, we used five individual samples from the acellular fraction of tuberculosis patients, along with three different pooled samples of the acellular fraction from additional TB patients, with each pool containing ten to twelve individual samples. For comparative purposes, we included one pooled sample of malignant PE (PE-C) from six patients, which encompassed cases of mesothelioma, lung carcinoma, or metastatic PE. Additionally, we used one pooled sample of parapneumonic PE (PE-PNI) from eight patients and one pooled sample of heart failure-related transudates (HF-PE) from five patients. The research was carried out in accordance with the Declaration of Helsinki (2013) of the World Medical Association. It was approved by the Institutional Review Board of Hospital F. J. Muñiz (approval number PRIISA.BA – 5334, renewed in December 2024) via the Buenos Aires Computerized Health Research Registry Platform. Written informed consent was obtained before sample collection.

### Sex as a biological variable

Both male and female individuals were included in the study, but due to the limited number of female participants, sex-based analyses could not be reliably performed.

### Bacterial strain

The Mtb laboratory strain, H37Rv, was grown at 37°C in Middlebrook 7H9 medium supplemented with 10% albumin-dextrose-catalase (both from Becton, Dickinson, New Jersey, USA) and 0.05% Tween-80 (Sigma-Aldrich, St. Louis, MO, USA). The red fluorescent protein (RFP)-expressing Mtb strain was gently provided by Dr. Fabiana Bigi (INTA, Castelar, Argentina).

### Preparation of PE pools

The PE were collected in heparin tubes and pools of the acellular fractions were prepared, as previously described^15^.

### Preparation of human monocyte-derived Macrophages

Monocytes were isolated from PBMC, as previously described^15^, and differentiated in the presence of human recombinant Granulocyte-Macrophage Colony-Stimulating Factor (GM-CSF, Peprotech, New York, USA) at 50 ng/ml. Cells were allowed to differentiate for five to seven days into M0 macrophages.

### Macrophage’s treatments

Macrophages were polarized toward M1 profile with IFNγ (10 ng/ml) plus LPS (10 ng/ml), exposed (or not) to either RvD5 (33 nM), vehicle (ethanol), or 20% v/v of PE during 24 h to generate M1 and treated-M1 cells, respectively. When indicated, macrophages were pre-incubated with GPR32-blocking antibodies (10 µg/mL; GTX71225, GeneTex) or their rabbit IgG isotype control (GeneTex) for 30 minutes before exposure to RvD5. Alternatively, M1 macrophages were generated only with IFNγ and either stimulated with Mtb γ-irradiated H37Rv strain (NR-49098, BEI Resources, USA) (equivalent to a multiplicity of infection “MOI” of 1:1) or infected with Mtb, and exposed (or not) to RvD5 (33 nM) for 24 h. Also, when specified, cells were exposed (or not) to DMOG (200 µM) during LPS or Mtb infection for 24 h.

In some experiments, M1 macrophages were treated with lipids extracted from PE. Briefly, lipids were isolated from PE, dried under a gentle nitrogen stream, and reconstituted in a small volume of methanol. This volume was normalized to the original volume of each pleural fluid sample to maintain the relative lipid abundances for subsequent cellular assays.

### Determination of lactate

Lactate production in the culture medium was measured using the Lactate Kit from Wiener (Argentina). The absorbance was read using a Biochrom Asys UVM 340 Microplate Reader and software.

### Lipid preparation and lipidomic analysis of PUFAs from TB-PE

Lipid extraction from TB-PE was based on the Bligh and Dyer protocol, as previously described^15^. PUFAs were then extracted on a solid phase as follows. 4 µL of the internal standards mixture (deuterium-labeled compounds) was added to the lipid extracts, which were resuspended in 300 µL of cold methanol. After centrifugation at 2000 g for 15 min at 4°C, the supernatants were transferred into 2 mL 96-well deep plates and diluted to 2 mL with H_2_O. Samples were then submitted to solid-phase extraction (SPE) using an OASIS HLB 96-well plate (30 mg/well, Waters), pretreated with MeOH (1 mL), and equilibrated with 10% MeOH (1 mL). After the sample application, the extraction plate was washed with 10% MeOH (1 mL). After drying under aspiration, lipid mediators were eluted with 1 mL of MeOH. Before LC-MS/MS analysis, samples were evaporated under nitrogen gas and reconstituted in 10 µL of MeOH.

The samples were analyzed at the MetaToul lipidomic platform (I2MC, INSERM 1048, Toulouse, France) as previously described^62^. Briefly, lipid mediators were separated on a ZorBAX SB-C18 column (2.1 mm, 50 mm, 1.8 µm) (Agilent Technologies) using an Agilent 1290 Infinity HPLC system (Agilent Technologies) coupled to an ESI-triple quadruple G6460 mass spectrometer (Agilent Technologies). Data was acquired in Multiple Reaction Monitoring (MRM) mode with optimized conditions (ion optics and collision energy). Peak detection, integration, and quantitative analysis were performed using Mass Hunter Quantitative Analysis software (Agilent Technologies) based on calibration lines constructed from commercially available eicosanoid standards (Cayman Chemicals).

### Determination of intracellular RvD5 abundances

Blood monocytes from healthy donors were stimulated with TB-PE (20% v/v) for 30 minutes. Cells were washed with ice-cold PBS (without Mg^2+^ and Ca^2+^) and scraped into ice-cold methanol/5 mM EGTA (2:1 v/v), snap-frozen in liquid nitrogen, and stored at-80°C until SPE and LC-MS analysis, as described above.

### SCENITH assays

SCENITH experiments were performed using the SCENITH kit, which contains all reagents and anti-puromycin antibodies (www.scenith.com). Briefly, PFMCs, PBMCs, and macrophages were treated for 40 minutes at 37°C in the presence of the indicated inhibitors of various metabolic pathways and puromycin. After incubation, puromycin was stained with fluorescently labeled anti-Puromycin monoclonal antibodies (clone R4743L-E8) conjugated to Alexa Fluor 647 or Alexa Fluor 488, and analyzed by flow cytometry. The impact of the various metabolic inhibitors was quantified, as previously described^27,28^.

### Measurement of cell respiration with Seahorse flux analyzer

Bioenergetics in human macrophages were determined using a Seahorse XFe24 analyzer. ATP production rates and relative contribution from glycolysis and oxidative phosphorylation (OXPHOS) were measured by the Seahorse XF Real-Time ATP Rate Assay kit. Macrophages (250,000 cells/well) were cultured in triplicate per condition. The assay was performed in XF Assay Modified DMEM. Three consecutive measurements were performed under basal conditions and after sequential addition of oligomycin (OM) and rotenone/antimycin (ROT/AA) (Agilent, USA). Extracellular acidification (ECAR) and oxygen consumption rates (OCR) were measured. Mitochondrial ATP production rate was determined by the decrease in the OCR after oligomycin addition. Conversely, the complete inhibition of mitochondrial respiration with rotenone plus antimycin A allows accounting for mitochondrial-associated acidification and, when combined with proton efflux rate (PER) data, calculation of the glycolytic ATP production rate. OCR and ECAR values were normalized by first performing brightfield imaging before the assay, then using ImageJ software to calculate the cellular area per condition, and finally importing these cellular area values into Agilent’s Wave software to divide the metabolic data by cell quantity, thereby accounting for variations in cell numbers between experiments and conditions.

Additionally, the Mito Stress Test was performed to measure mitochondrial oxidative phosphorylation rate. Basal respiration was calculated as the last measurement before the addition of OM minus the non-mitochondrial respiration, which is the minimum rate measurement after the addition of ROT/AA. Estimated ATP production is the last measurement before OM addition minus the minimum rate after OM. Maximal respiration rate (Max) was defined as the OCR after addition of OM and FCCP at 2µM (for 2.5 minutes) minus the non-mitochondrial OCR. Spare respiratory capacity (SRC) was defined as the difference between maximum and basal respiration. Three consecutive measurements were performed under basal conditions and after the sequential addition of the following electron transport chain inhibitors.

### RNA-Single Cell Sequencing Analysis

Previously published single cell transcriptomic data available in Genome Sequence Archive in the BIG Data Center, Beijing Institute of Genomics, Chinese Academy of Sciences, under accession numbers HRA000910 and HRA00036^63^, were used to evaluate the expression of enzymes involved in SPMs’ biosynthesis (ALOX5, ALOX12, ALOX 15 and ALOX15B), including five samples of PBMC and six samples of PFMC from TB patients. The Seurat package (version 4.3.0.1) in RStudio was used to read the pre-processed files. The filtering process involved discarding cells with fewer than 200 or more than 5000 total genes, fewer than 200 or more than 20,000 total counts, and mitochondrial read levels above 10%. Expression levels were then normalized using the log transformation (scale factor = 10,000), based on the 2,000 most variable genes. The samples were integrated using the reciprocal PCA (RPCA) algorithm. The different cell clusters were identified using the shared nearest neighbor (SNN) modularity optimization algorithm, and dimensionality reduction was performed using the UMAP (Uniform Manifold Approximation and Projection) algorithm. Individual cells were annotated using the SingleR package version 2.2.0 and the HumanPrimaryCellAtlasData database (label.fine version).

### Intracellular expression of HIF1α

Macrophages were permeabilized with triton 0.3% in FACS buffer and incubated with PE-anti-HIF1α (clone 546-16, Biolegend) for 30 minutes. After that, cells were analyzed by flow cytometry using the FACSCalibur cytometer (BD Biosciences, San Jose, CA, USA). The macrophage population was gated according to its Forward and Size Scatter properties. The median fluorescence intensity (MFI) was analyzed using FCS Express V3 software (*De Novo* Software, Los Angeles, CA, USA).

### GPR32 surface expression

To monitor GPR32 expression in CD14^+^ cells from pleural effusion and blood paired samples of TB vs. non-TB patients, cells were incubated with a primary anti-GPR32 antibody (GTX71225; GeneTex) for 40 minutes at 4°C. Next, the cells were incubated with a secondary antibody conjugated to Alexa Fluor 488 (anti-Rabbit IgG, 111-546-144) to label anti-GPR32. After an anti-CD14-PerCP/Cy7 (clone HCD14, Biolegend) was employed, the cell viability marker Zombie Violet (Biolegend) was used.

### Fluorescence microscopy

Fluorescence microscopy images of fixed IFNγ-activated (M1) macrophages exposed (or not) to RvD5 that were first infected with Mtb-RFP (MOI 1-3:1) for 24 h and then fixed. Cell nuclei were labeled with DAPI, and the autophagy marker LC3 protein (anti-LCE3B; dilution 1:300; #NB100-2220) was identified by indirect immunofluorescence using Alexa Fluor 488 conjugated antibodies, whereas the late endosome marker, LAMP2, was identified by indirect immunofluorescence using a specific conjugated antibody (anti-LAMP2 Alexa Fluor 633; dilution 1:200; #A21126). Samples were imaged on a FluoView FV1000 confocal microscope (Olympus, Tokyo, Japan) equipped with a Plapon 60X/NA1.42 objective and on the Zeiss LSM 800 confocal microscope (Zeiss, Oberkochen, Germany).

### IL-1β determination

The amount of human IL-1β was measured by ELISA, according to the manufacturer’s instructions (ELISA MAX Deluxe Kits from Biolegend). The detection limit was 15.6 pg/ml.

### Infection of human macrophages with Mtb

Infections were performed in the biosafety level 3 laboratories at the *Unidad Operativa Centro de Contención Biológica (UOCCB), ANLIS-MALBRAN* (in 2022) and at *Instituto de Investigaciones Biomédicas en Retrovirus y SIDA (INBIRS)-CONICET-UBA* (in 2023-24), according to the biosafety institutional guidelines (both institutes located in Buenos Aires). Macrophages were infected with Mtb H37Rv strain at a MOI of 0.5-3:1 during 1 h at 37°C. Next, extracellular bacteria were removed gently by washing with pre-warmed PBS, and cells were cultured in RPMI-1640 medium supplemented with 10% FBS and gentamicin (50 μg/ml) for 4, 24 and 72h. At different time points, macrophages were lysed in 0.1% SDS and neutralized with 20% BSA in Middlebrook 7H9 broth. Serial dilutions of the lysates were plated in triplicate, onto 7H11-Oleic Albumin Dextrose Catalase (OADC, Becton Dickinson) agar medium for CFU scoring at 21 days later.

### Western blot

Macrophages were stimulated with IFNγ in the presence or not of RvD5, or left unstimulated, for 24 h. Cells were then infected with Mtb H37Rv for 6 h and lysed using RIPA Lysis Buffer System (Santa Cruz, CAT #sc-24948). Lysates were cleared by centrifugation at 15,000 rpm for 5 minutes at 4°C. Protein concentrations were determined using the Pierce BCA Protein Assay Kit (Thermo Fisher, CAT #23227). Equal amounts of protein (25 µg per lane) were resolved on a 15% SDS-PAGE gel. Separated proteins were transferred onto a PVDF membrane (Roche, CAT #03010040001) at 350 mA for 1 h. The membrane was blocked with 5% skimmed milk for 1 h at room temperature to prevent non-specific binding. Membranes were incubated overnight at 4°C with a primary anti-human LC3B antibody (1:1000 dilution, Novus Biologicals, CAT #NB100-2220). After extensive washing, blots were incubated with a horseradish peroxidase (HRP)-conjugated goat anti-rabbit IgG secondary antibody (1:10000 dilution, Jackson ImmunoResearch, CAT #111-035-003) for 1 h at room temperature. Immunoreactivity was detected using SuperSignal™ West Pico PLUS Chemiluminescent Substrate (Thermo Fisher). Protein bands were visualized on Kodak Medical X-Ray General Purpose Film. For loading control, membranes were extensively washed and reprobed with anti-beta actin antibody (1:1000 dilution, BD Mouse Anti-Actin Ab-5; CAT #612657), followed by HRP-conjugated goat anti-mouse IgG antibody (1:10000 dilution). Band intensities were quantified by densitometric analysis using ImageJ software.

### Quantification and Statistical Analysis

Data are presented as scatter plots where each symbol represents one independent donor or biological sample, with means and standard error of the mean (SEM) shown unless otherwise indicated. Each experiment corresponds to a separate donor. Statistical tests were selected based on experimental design and data distribution. Comparisons between two unpaired groups used the Mann–Whitney test or unpaired t-test; comparisons between two paired groups used the Wilcoxon signed-rank test or paired t-test. Comparisons among three or more groups used one-way or two-way ANOVA followed by post hoc tests: Dunnett’s test for comparisons against a single control; Tukey’s test for comparisons among all groups; Holm–Sidak test for specified comparisons in multi-factor designs; or the Benjamini–Krieger–Yekutieli two-stage linear step-up procedure to control the false discovery rate. Categorical data were analyzed with Fisher’s exact test, correlation analyses used Spearman’s rank correlation coefficient, and specialized designs employed nested t-tests for hierarchical data or mixed-effects models for repeated measures. Specific tests for each figure are detailed in the corresponding legends. A p-value < 0.05 was considered significant. Analyses were performed using GraphPad Prism (version 8.0.1) or R.

## Data availability statement

All data supporting the findings of this study are available within the article and its Supplementary Information. The lipidomics datasets generated during this study have been deposited in the Metabolights repository under accession code MTBLS12926 and will be publicly available as of 2026-04-07.

## Author contributions

Conceptualization: CV, GLV, and LB. Methodology: JB and LB. Investigation: JB, MMaio, LF, EB, NF, SM, JLMF, AM, FS, FF, TMG, MMatteo, EL, CV, GLV, and LB. Resources: AJE, XA, MSC, DP, MO, RA, RKN, ON, CV, GLV, and LB. Formal Analysis: JB, MML, EL, and LB. Writing: JB, MML, EL, ON, CV, GLV, and LB. Visualization: JB, FF, and LB. Supervision: EL, CV, GLV, and LB. Project Administration: LB. Funding acquisition: RA, ON, and LB. Corresponding author: LB is responsible for ownership and responsibility that are inherent to aspects of this study.

## Acknowledgements

We thank the staff of the Regional Center of Hemotherapy of the Garrahan Hospital (Buenos Aires). We thank MetaToul (Toulouse metabolomics & fluxomics facilities, https://mth-metatoul.com/), which is part of the French National Infrastructure for Metabolomics and Fluxomics, MetaboHUB-ANR-11-INBS-0010 (doi.org/10.15454/VRJK-KY76). We also thank S. Fried and A. Lucas from the We-Met platform at I2MC, in Toulouse, France. We also thank Life Science Editors for editing services (www.lifescienceeditors.com). This work was supported by the Argentinean National Agency of Promotion of Science and Technology (PICT-2019-01044 and PICT-2020-00501 to LB); the Argentinean National Council of Scientific and Technical Investigations (CONICET, PIP 11220200100299CO to LB); the ICGEB CRP/ARG23-02 Research Grant to LB; Fondation Bettencourt Schueller (EXPORE-TB grant to ON); MSDAVENIR (the FIGHT-TB grant to ON); and the French ANR JCJC-Epic-SCENITH ANR-20-CE14-0028 and CoPoC Inserm-transfert MAT-PI-17493-A-04 to RA.

