## Supplementary Figures for "Lipid-mediated GPR32 signaling reprograms macrophage metabolism to impair anti-tuberculous immunity"

**SFigure 1**

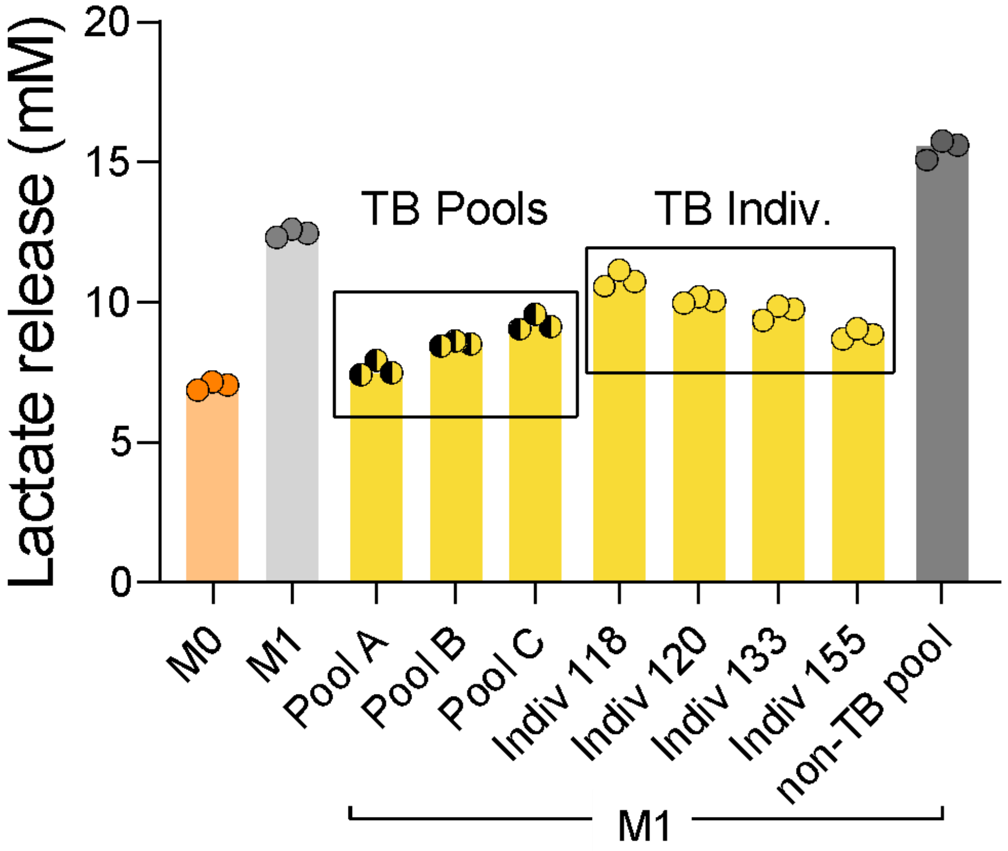

Lactate release (mM)
20
15
10
5
0
TB Pools
TB Indiv.
M0
M1
Pool A
Pool B
Pool C
Indiv 118
Indiv 120
Indiv 133
Indiv 155
non-TB pool
M1

### SFigure 2

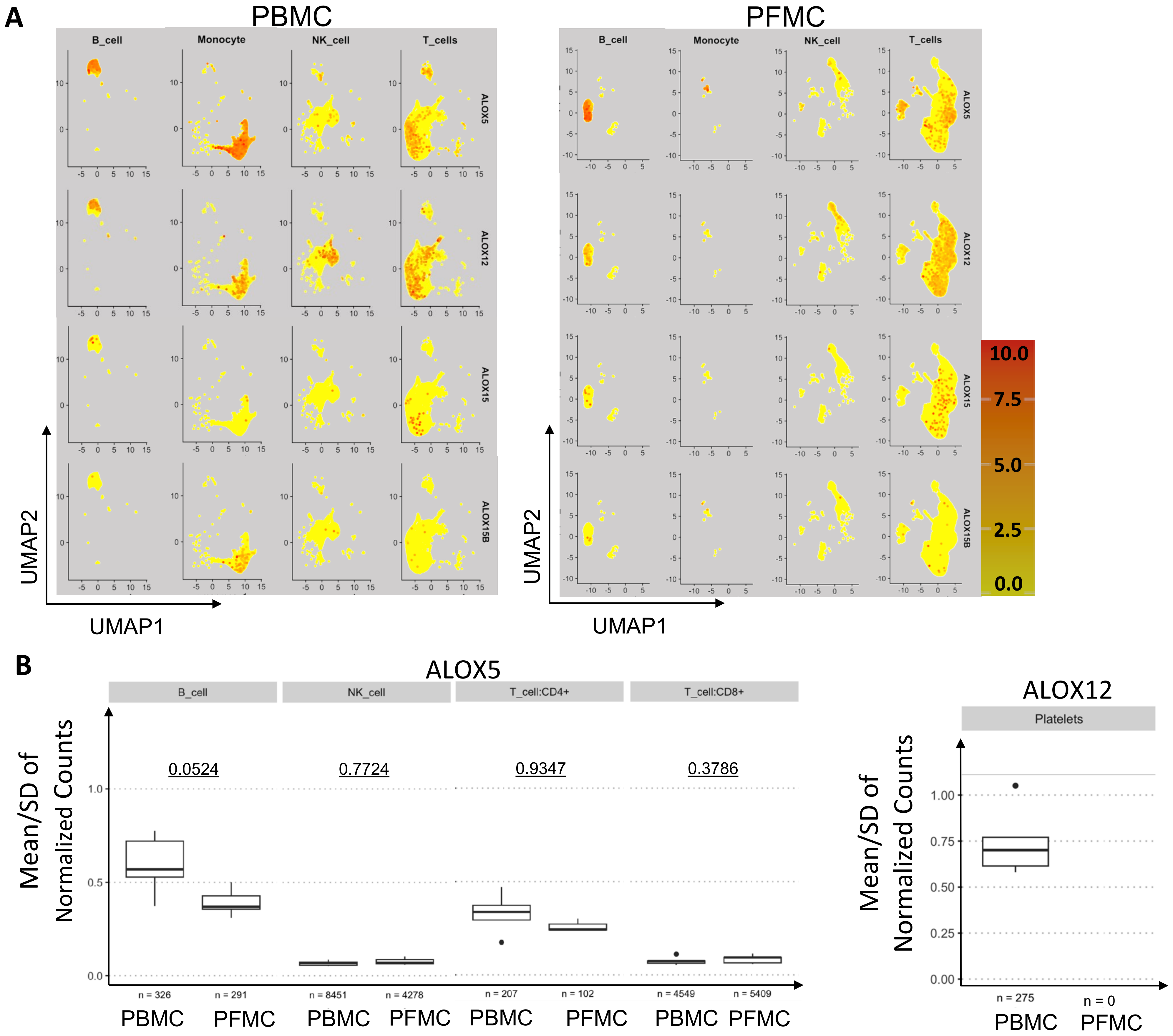
A
PBMC
PFMC
B_cell
Monocyte
NK_cell
T_cells
ALOX5
ALOX12
ALOX15
ALOX15B
UMAP2
UMAP1
10.0
7.5
5.0
2.5
0.0
B
ALOX5
B_cell
NK_cell
T_cell:CD4+
T_cell:CD8+
0.0524
0.7724
0.9347
0.3786
Mean/SD of Normalized Counts
n = 326
n = 291
n = 8451
n = 4278
n = 207
n = 102
n = 4549
n = 5409
PBMC
PFMC
ALOX12
Platelets
n = 275
n = 0

### SFigure 3

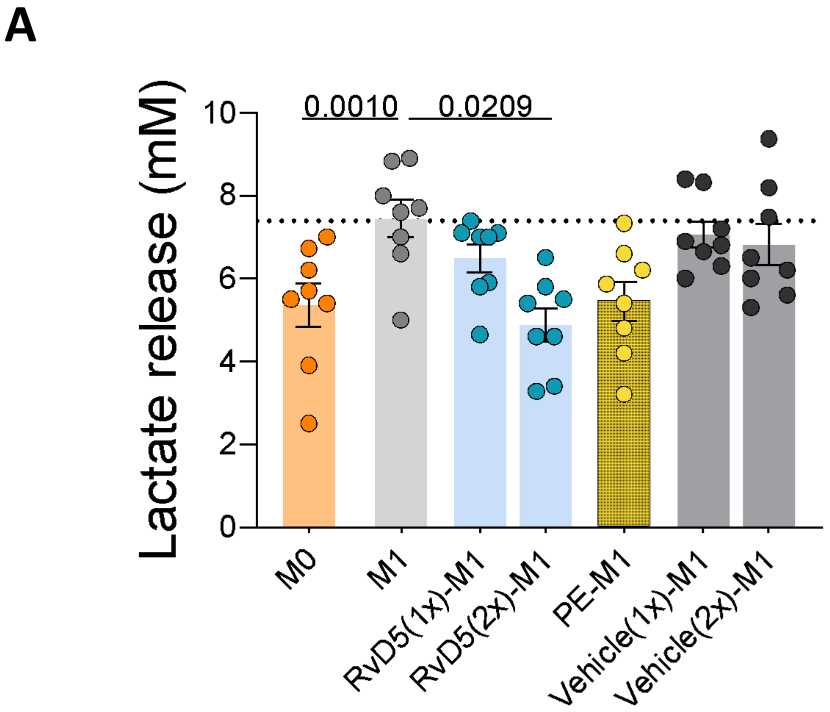
A
Lactate release (mM)
0.0010
0.0209
10
8
6
4
2
0
M0
M1
RvD5(1x)-M1
RvD5(2x)-M1
PE-M1
Vehicle(1x)-M1
Vehicle(2x)-M1

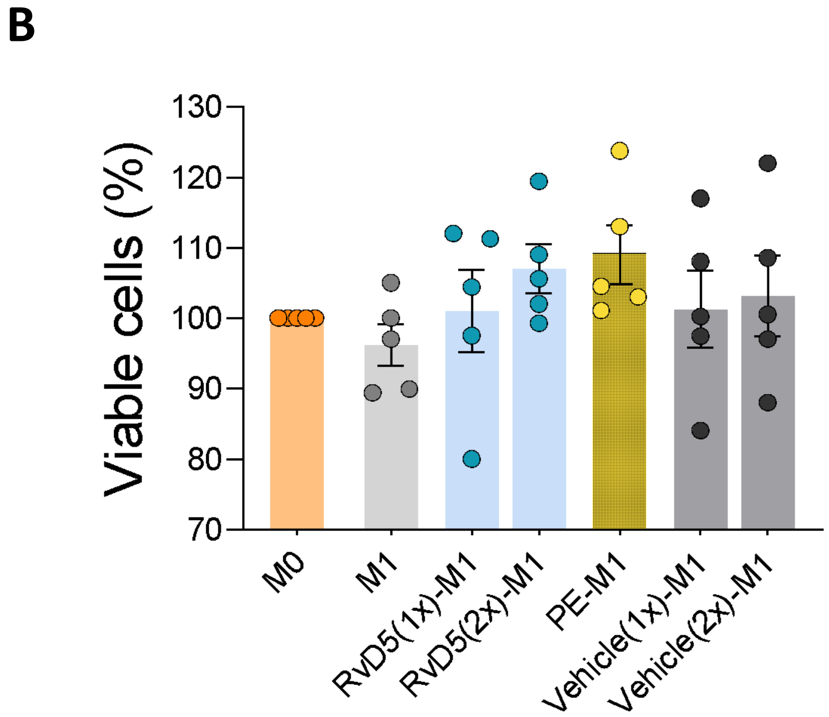
B
Viable cells (%)
130
120
110
100
90
80
70
M0
M1
RvD5(1x)-M1
RvD5(2x)-M1
PE-M1
Vehicle(1x)-M1
Vehicle(2x)-M1

### SFigure 4

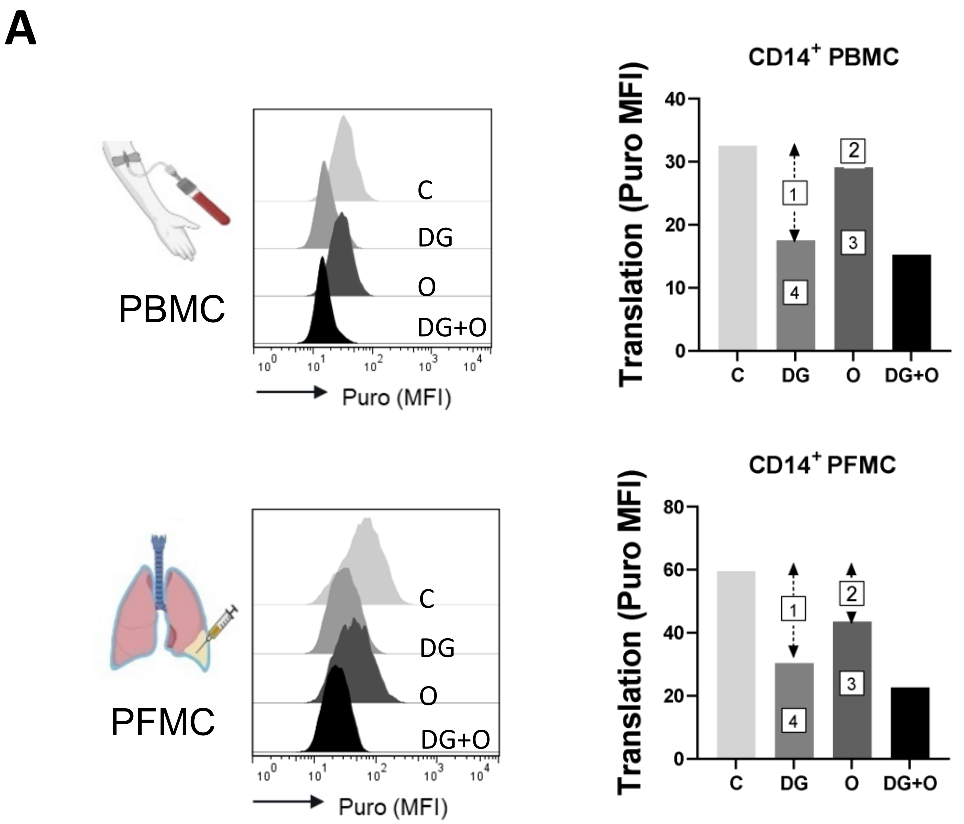
A
PBMC
C
DG
O
DG+O
Puro (MFI)
CD14+ PBMC
Translation (Puro MFI)
C
DG
O
DG+O
PFMC
CD14+ PFMC

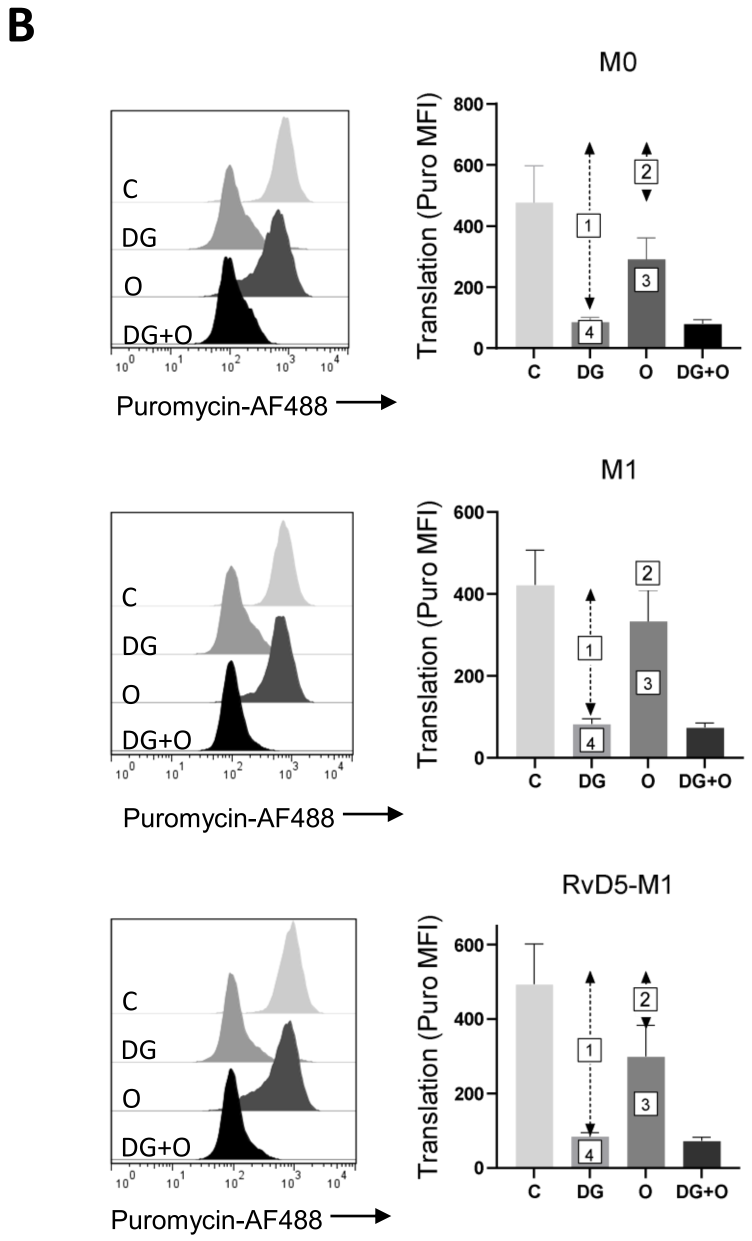
B
M0
M1
RvD5-M1
C
DG
O
DG+O
Puromycin-AF488
Translation (Puro MFI)

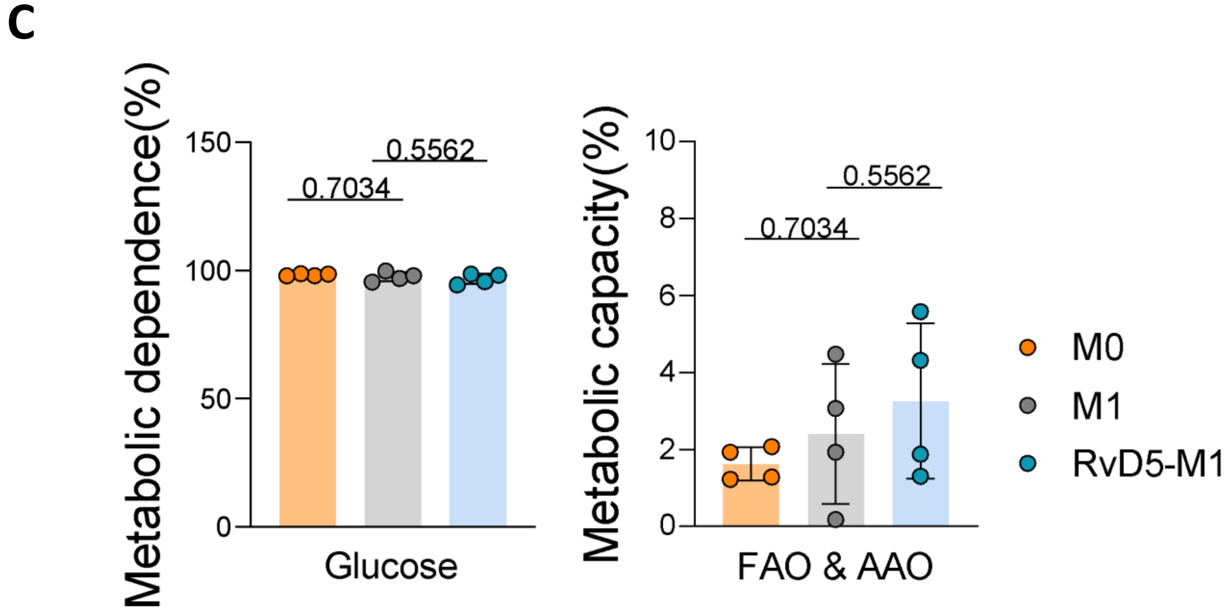
C
Metabolic dependence(%)
0.7034
0.5562
Glucose
Metabolic capacity(%)
FAO & AAO
M0
M1
RvD5-M1

### SFigure 5

**A**

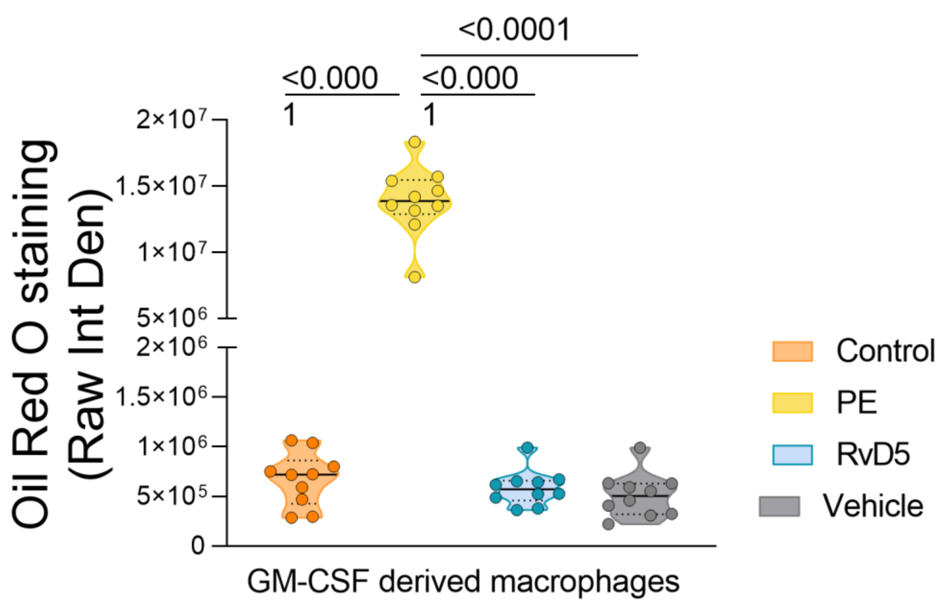

<0.0001
<0.000
1
<0.000
1
2×10⁷
1.5×10⁷
1×10⁷
5×10⁶
2×10⁶
1.5×10⁶
1×10⁶
5×10⁵
0
Oil Red O staining
(Raw Int Den)
GM-CSF derived macrophages
Control
PE
RvD5
Vehicle


**B**

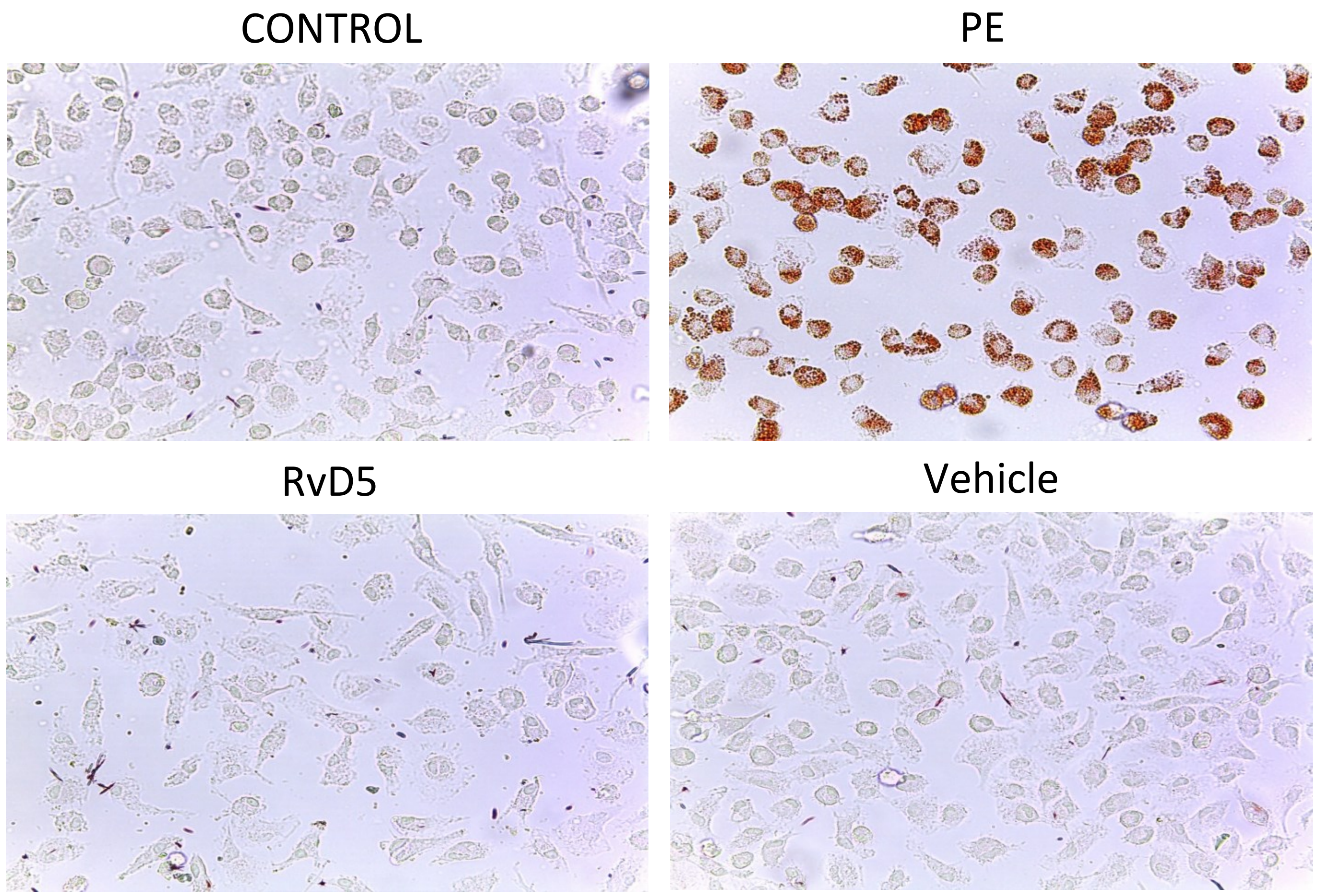

CONTROL
PE
RvD5
Vehicle

### SFigure 6

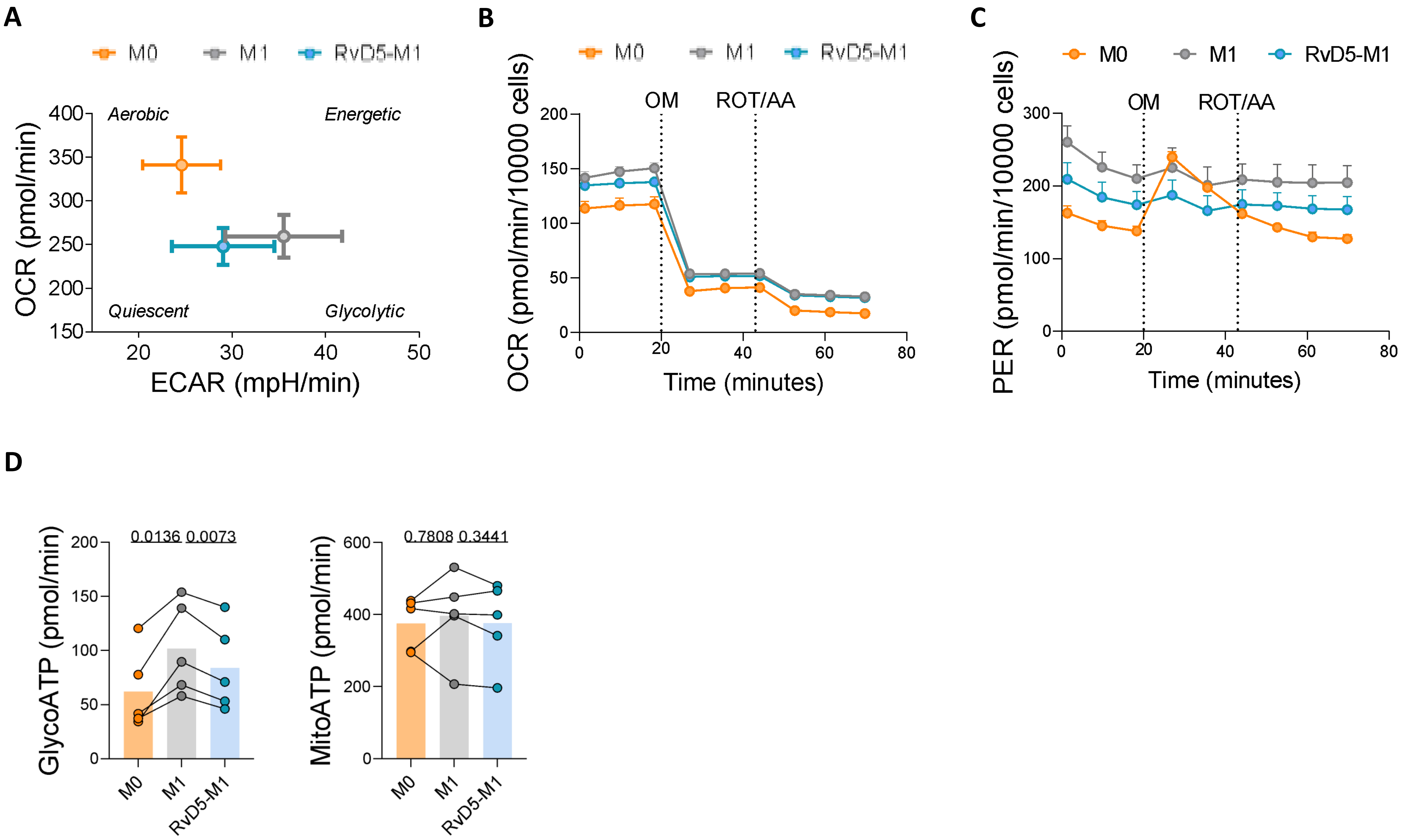
A
M0
M1
RvD5-M1
OCR (pmol/min)
ECAR (mpH/min)
Aerobic
Energetic
Quiescent
Glycolytic
400
350
300
250
200
150
20
30
40
50
B
OCR (pmol/min/10000 cells)
OM
ROT/AA
Time (minutes)
200
150
100
50
0
0
20
40
60
80
C
PER (pmol/min/10000 cells)
300
200
100
0
D
GlycoATP (pmol/min)
0.0136
0.0073
MitoATP (pmol/min)
0.7808
0.3441
600
400
200
0

### SFigure 7

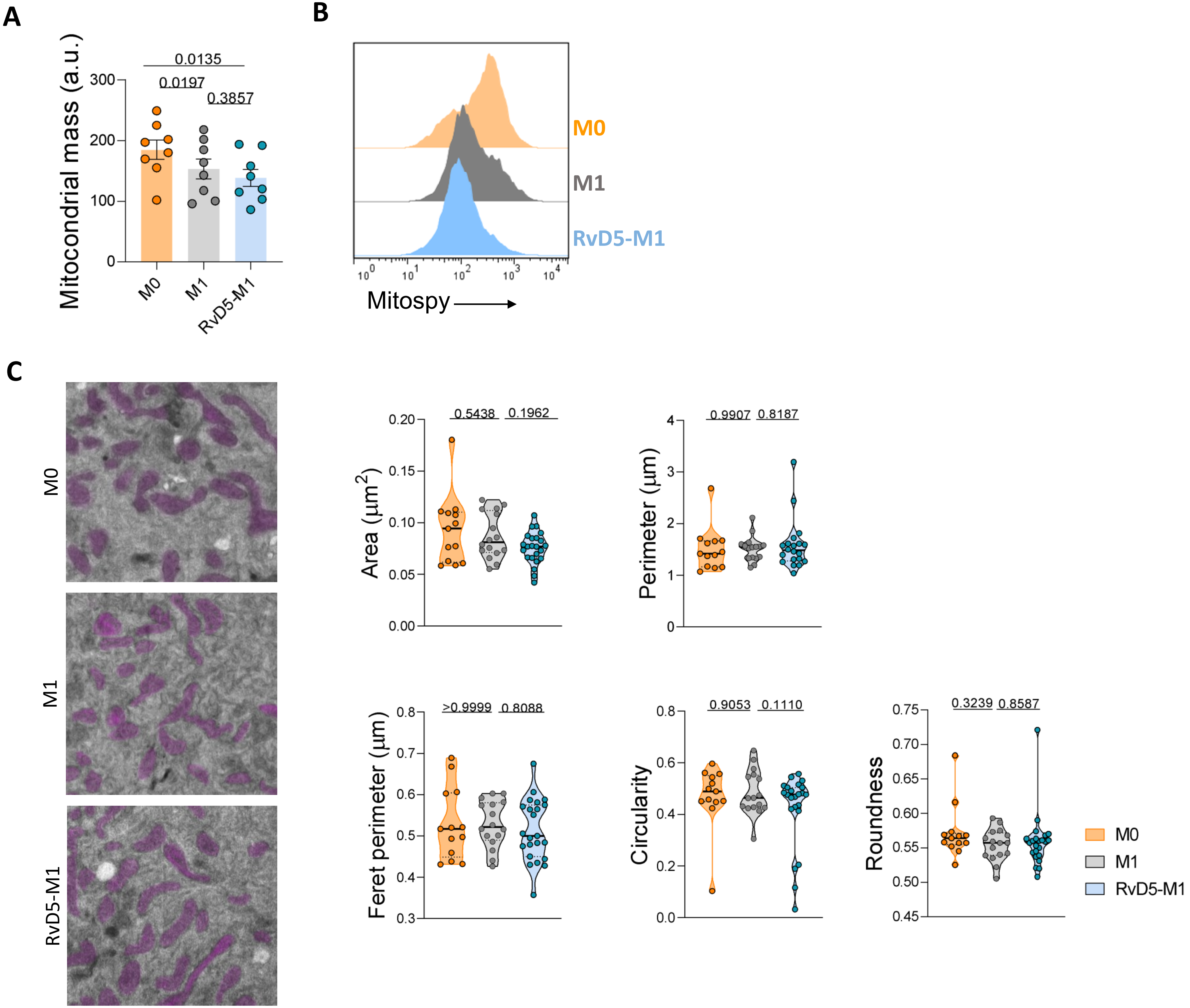

A
Mitocondrial mass (a.u.)
0.0135
0.0197
0.3857
300
200
100
0
M0
M1
RvD5-M1
B
M0
M1
RvD5-M1
Mitospy
C
M0
M1
RvD5-M1
Area (μm2)
0.5438
0.1962
Perimeter (μm)
0.9907
0.8187
Feret perimeter (μm)
>0.9999
0.8088
Circularity
0.9053
0.1110
Roundness
0.3239
0.8587
M0
M1
RvD5-M1

### SFigure 8

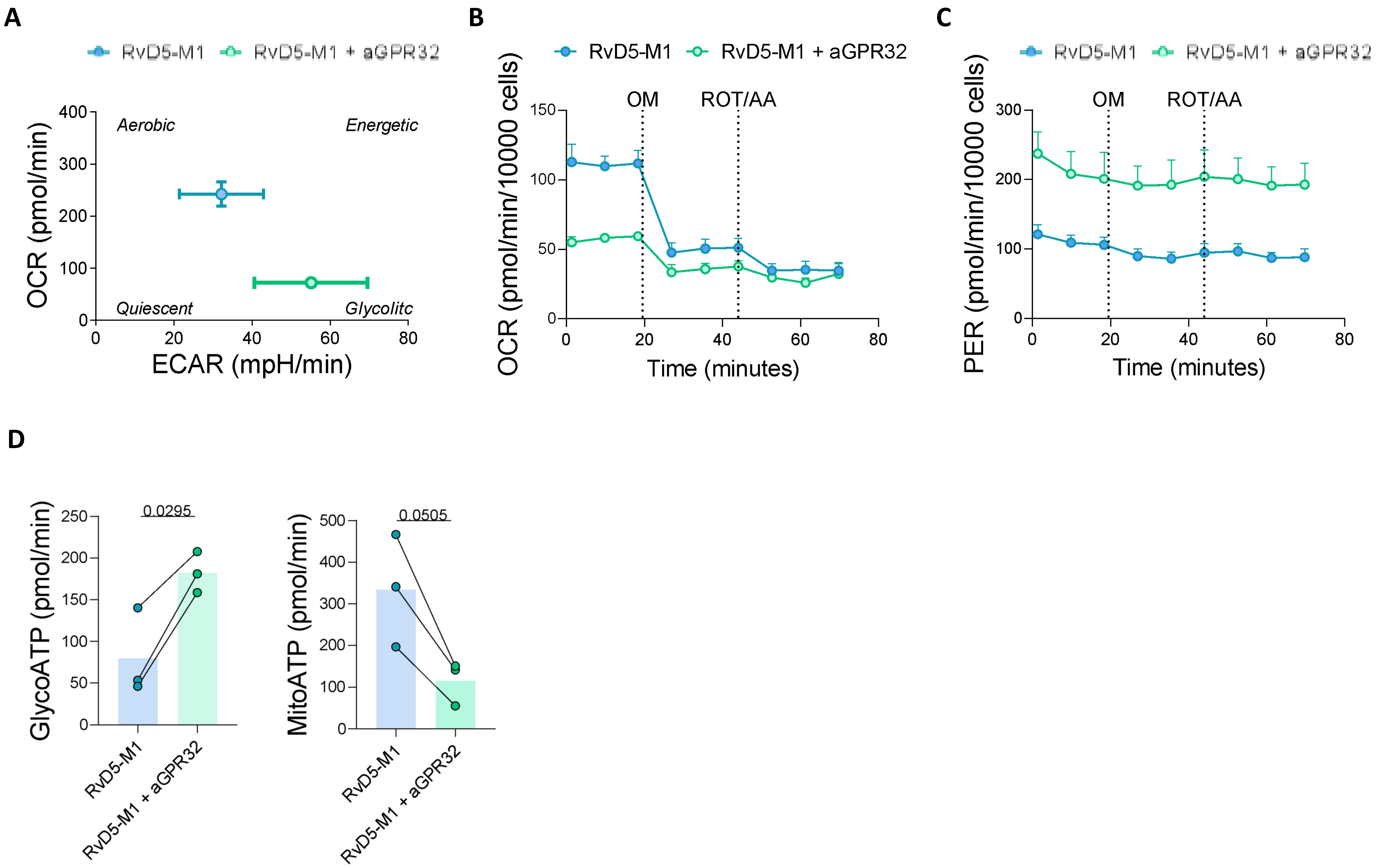

A
RvD5-M1
RvD5-M1 + aGPR32
OCR (pmol/min)
ECAR (mpH/min)
Aerobic
Energetic
Quiescent
Glycolitc
B
RvD5-M1
RvD5-M1 + aGPR32
OCR (pmol/min/10000 cells)
OM
ROT/AA
Time (minutes)
C
RvD5-M1
RvD5-M1 + aGPR32
PER (pmol/min/10000 cells)
OM
ROT/AA
Time (minutes)
D
GlycoATP (pmol/min)
0.0295
RvD5-M1
RvD5-M1 + aGPR32
MitoATP (pmol/min)
0.0505
RvD5-M1
RvD5-M1 + aGPR32

### SFigure 9

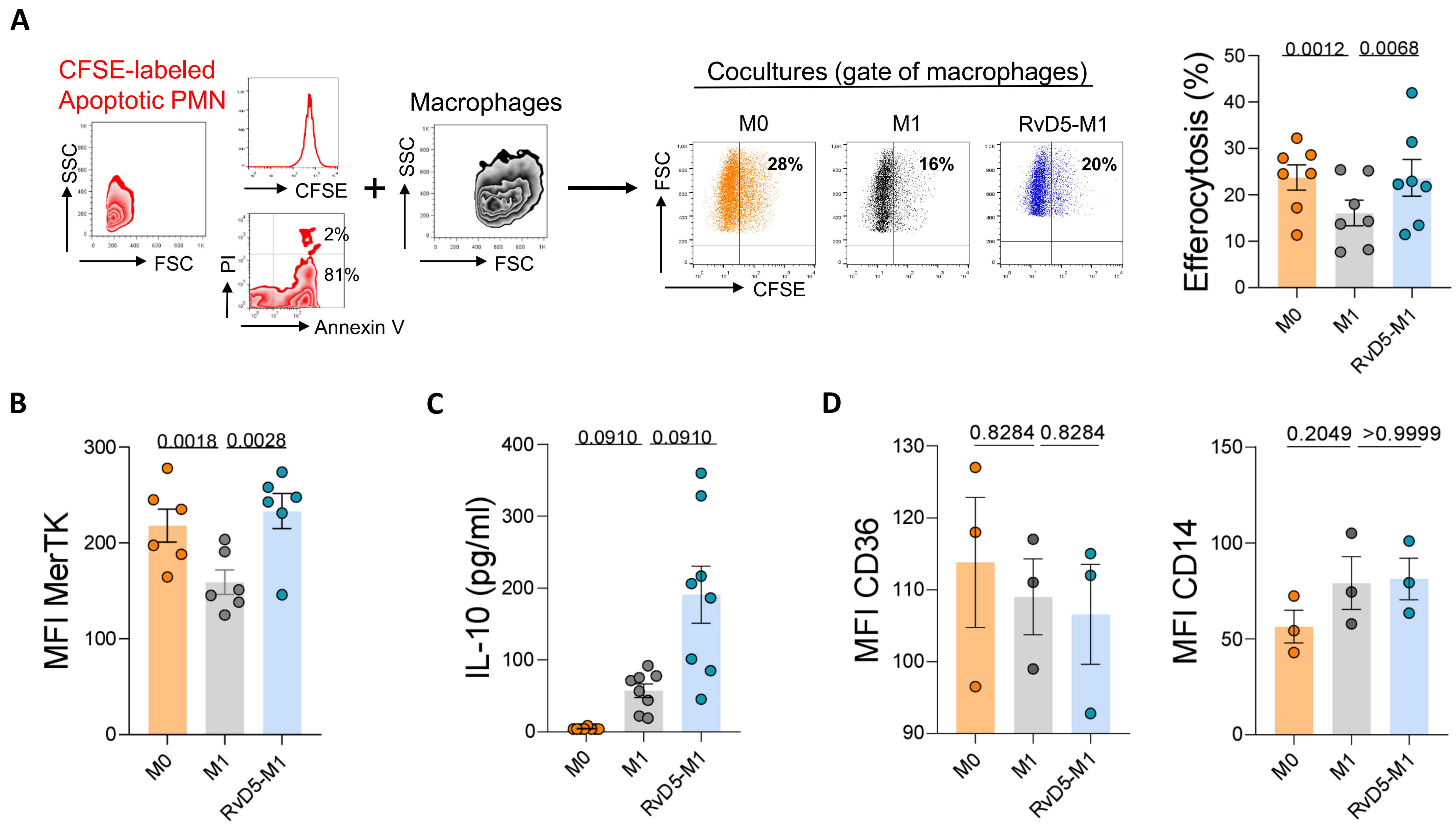
A
CFSE-labeled
Apoptotic PMN
SSC
FSC
CFSE
PI
Annexin V
2%
81%
Macrophages
SSC
FSC
Cocultures (gate of macrophages)
M0
M1
RvD5-M1
FSC
CFSE
28%
16%
20%
Efferocytosis (%)
0.0012
0.0068
M0
M1
RvD5-M1
B
MFI MerTK
0.0018
0.0028
C
IL-10 (pg/ml)
0.0910
0.0910
D
MFI CD36
0.8284
0.8284
MFI CD14
0.2049
>0.9999

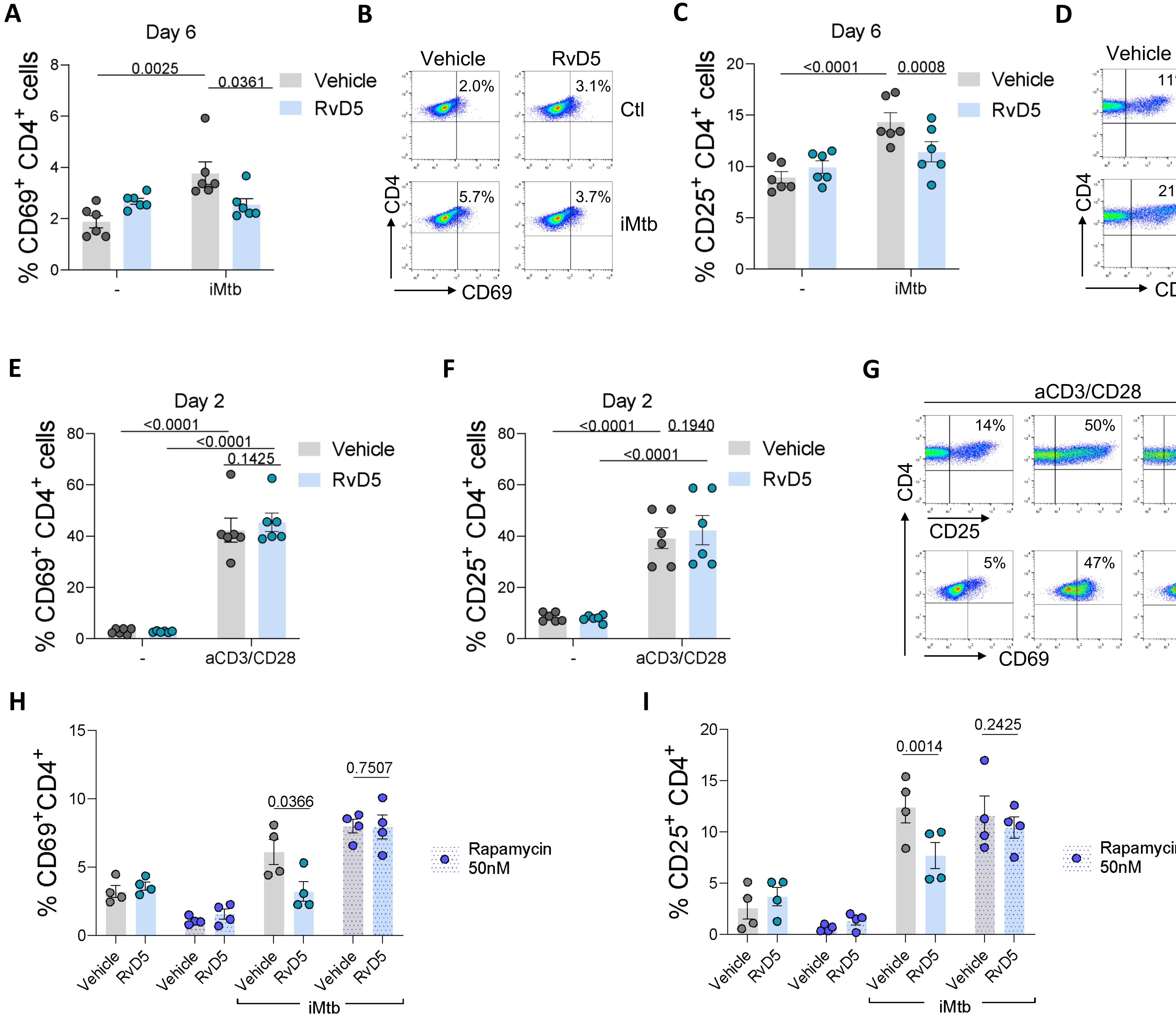

A
Day 6
% CD69+ CD4+ cells
0.0025
0.0361
Vehicle
RvD5
-
iMtb
B
Vehicle
RvD5
2.0%
3.1%
Ctl
5.7%
3.7%
iMtb
CD4
CD69
C
Day 6
% CD25+ CD4+ cells
<0.0001
0.0008
Vehicle
RvD5
-
iMtb
D
Vehicle
RvD5
11%
12%
Ctl
21%
17%
iMtb
CD4
CD25
E
Day 2
% CD69+ CD4+ cells
<0.0001
<0.0001
0.1425
Vehicle
RvD5
-
aCD3/CD28
F
Day 2
% CD25+ CD4+ cells
<0.0001
0.1940
<0.0001
Vehicle
RvD5
-
aCD3/CD28
G
aCD3/CD28
14%
50%
52%
CD4
CD25
5%
47%
51%
CD69
H
% CD69+CD4+
0.0366
0.7507
Rapamycin 50nM
Vehicle
RvD5
iMtb
I
% CD25+ CD4+
0.0014
0.2425
Rapamycin 50nM
Vehicle
RvD5
iMtb


**SFigure 10**

SFigure 11

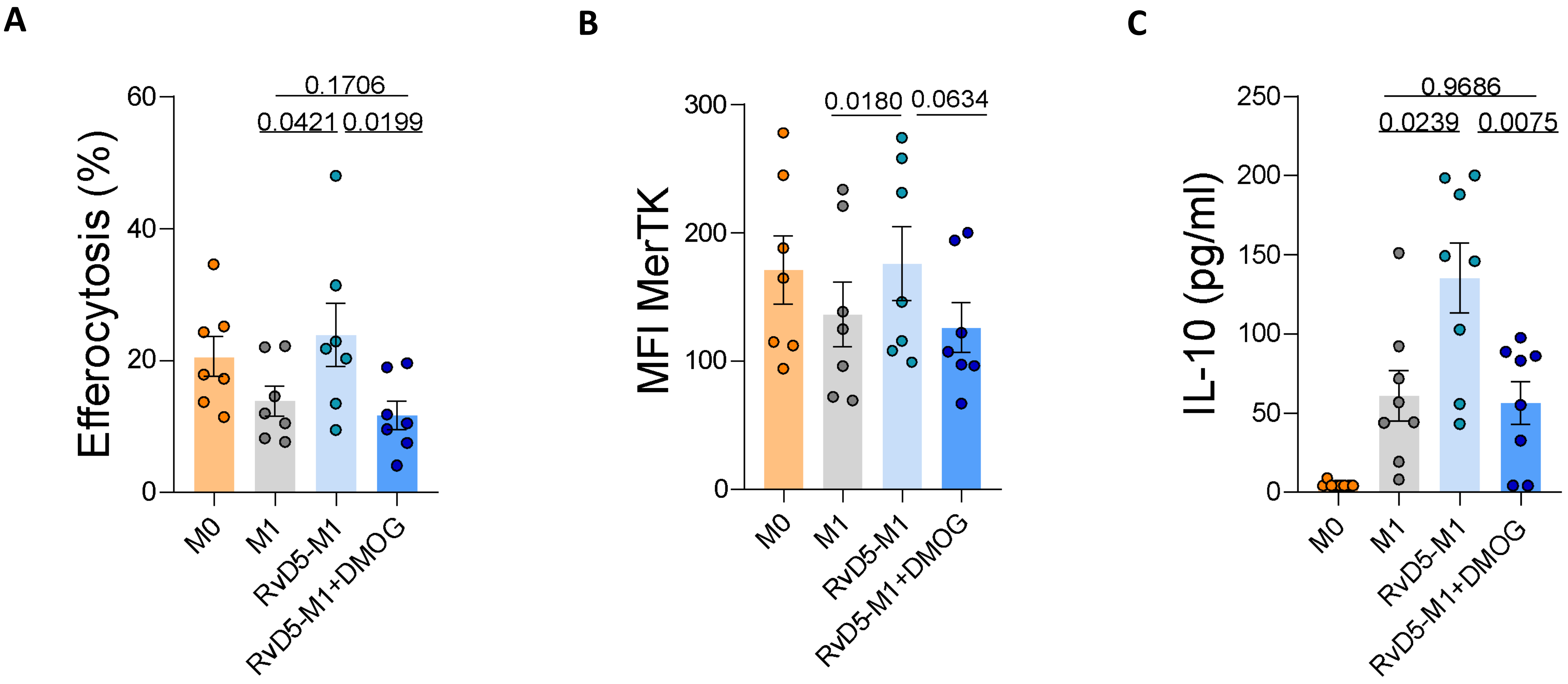
A
Efferocytosis (%)
60
40
20
0
0.1706
0.0421
0.0199
M0
M1
RvD5-M1
RvD5-M1+DMOG
B
MFI MerTK
300
200
100
0
0.0180
0.0634
M0
M1
RvD5-M1
RvD5-M1+DMOG
C
IL-10 (pg/ml)
250
200
150
100
50
0
0.9686
0.0239
0.0075
M0
M1
RvD5-M1
RvD5-M1+DMOG
