## Supplementary Tables I and II for "Lipid-mediated GPR32 signaling reprograms macrophage metabolism to impair anti-tuberculous immunity"

**Supplementary Table I** - Relative lactate release from M1 macrophages exposed to pooled vs. individual TB‑PE samples

| Sample | Replicate 1 | Replicate 2 | Replicate 3 | Mean ± SD |
| --- | --- | --- | --- | --- |
| Pool A | 0.6362 | 0.5933 | 0.6020 | 0.6105 ± 0.0227 |
| Pool B | 0.6821 | 0.6835 | 0.6842 | 0.6833 ± 0.0011 |
| Pool C | 0.7254 | 0.7257 | 0.7771 | 0.7427 ± 0.0296 |
| Ind 118 | 0.8622 | 0.8397 | 0.9037 | 0.8685 ± 0.0322 |
| Ind 120 | 0.8063 | 0.8093 | 0.8094 | 0.8083 ± 0.0018 |
| Ind 133 | 0.7826 | 0.7830 | 0.7586 | 0.7747 ± 0.0140 |
| Ind 155 | 0.7102 | 0.6897 | 0.7336 | 0.7112 ± 0.0220 |

*Macrophages were treated for 24 h with 20% v/v of the indicated pleural effusion samples. The relative response was calculated for each replicate as (lactate value) / (lactate of M1 macrophages in the same replicate). TB-PE pools A, B, and C each contained four distinct individual TB-PE samples (12 unique samples total). Individuals 118, 120, 133, and 155 were independent donors not included in any pool. All conditions were tested in technical triplicate.*

**Supplementary Table II -** Statistical comparison of relative lactate release between pooled and individual TB‑PE samples.

| Group | N | Mean Relative Response | Standard Deviation (SD) | Coefficient of Variation (CV) |
| --- | --- | --- | --- | --- |
| **Pools** | 3 | 0.6788 | 0.0662 | 9.76% |
| **Individuals** | 4 | 0.7907 | 0.0835 | 10.56% |

*M1 macrophages were treated with either three TB‑PE pools (each containing four distinct individual samples; n=3) or four individual TB‑PE samples from independent donors not included in any pool (n=4). The relative response was calculated for each technical replicate as (lactate value) / (lactate of M1 macrophages in the same replicate). An unpaired two-tailed t-test assuming equal variances was used to compare the mean relative responses between the two groups.*
