## Supplementary Legends for "Lipid-mediated GPR32 signaling reprograms macrophage metabolism to impair anti-tuberculous immunity"

[1] Instituto de Investigaciones Biomédicas en Retrovirus y SIDA (INBIRS). Facultad de Medicina. Universidad de Buenos Aires- Consejo Nacional de Investigaciones Científicas y Técnicas (CONICET), Buenos Aires, Argentina

[2] Instituto de Medicina Experimental (IMEX)-CONICET, Academia Nacional de Medicina, Buenos Aires, Argentina

[3] International Associated Laboratory (LIA) CNRS IM-TB/HIV (1167), Buenos Aires, Argentina / International Research Project Toulouse, France

[4] Univ Toulouse, CNRS, IPBS, Toulouse, France

[5] Instituto de Estudios Inmunológicos y Fisiopatológicos - CONICET - Universidad Nacional de La Plata, La Plata, Argentina

[6] Instituto Prof. Dr. Raúl Vaccarezza; Hospital de Infecciosas Dr. F.J. Muñiz, Buenos Aires, Argentina

[7] Aix-Marseille University, CNRS, INSERM, CIML, Centre d'Immunologie de Marseille-Luminy, Turing Centre for Living Systems, Marseille, France

[8] Translational Health Group, International Center of Genetic Engineering and Biotechnology, New Delhi, India

### Supplementary figure legends

**Figure S1. Validation of pooled TB-PE samples as biological proxies for individual TB-PE samples.** Lactate release from unstimulated (M0) or LPS/IFN-γ-activated (M1) macrophages following 24 h treatment with 20% v/v of the following pleural effusion samples: three pools of tuberculous pleural effusion (TB-PE), each pool comprising four distinct individual TB-PE samples (12 unique samples total across the three pools); four individual TB-PE samples from independent donors not included in any pool; and one non-TB pool. All conditions were tested in technical triplicates. This experiment was designed to assess whether pooled samples recapitulate the biological variability and average response of individual TB-PE samples, thereby justifying their inclusion in subsequent lipidomic analyses (see Supplementary Tables I and II for statistical comparison).

**Figure S2. Single-Cell RNA-seq profiles of Enzymes Involved in Specialized Pro-resolving lipid Mediators Biosynthesis in Mononuclear Cells from Pleural Effusions. (A)** 2D UMAP plots display mononuclear cells color-coded according to the expression levels of selected genes (ALOX5, ALOX12, ALOX15, and ALOX15B) across different cell types (B cells, monocytes, NK and T cells). *Left panel:* Profiles from blood mononuclear cells (PBMC, n=5). *Right panel:* Profiles from pleural mononuclear cells (PFMC, n=6). **(B)** Box-and-whisker plots show the mean-to-variability ratio (μ/σ) of normalized ALOX5 expression across B cell, NK cell, and $CD4^+$ and $CD8^+$ T cell lineages from TB patients, comparing peripheral blood (PBMC, n=5) and pleural fluid (PFMC, n=6). A final panel displays the same μ/σ ratio for normalized ALOX12 expression in platelets as a positive control. The total number of cells in each group is indicated on the x-axis. For each lineage and compartment, the ratio was calculated per patient. Boxes indicate the interquartile range (IQR) of these patient-level ratios, whiskers extend to 1.5 × IQR, and center lines denote the median. Statistical comparisons between PBMC and PFMC for each lineage were performed using a nested t-test.

**Figure S3. Impact of the lipid mediator RvD5 on lactate release. (A)** Lactate release from unstimulated (M0) or LPS- and IFNγ-activated (M1) macrophages treated for 24 h with RvD5 (17 and 33 nM) or vehicle (ethanol). Supernatant lactate was measured using an enzymatic colorimetric assay (n=8). Data was analyzed by one-way ANOVA followed by Sidak´s multiple comparison test. **(B)** Cell viability assessed via MTT assay. Results are expressed as the percentage of viable cells relative to M0 controls.

**Figure S4. Impact of RvD5 on Macrophage Metabolism.** (A) Metabolic profiling of $CD14^+$ cells from blood or pleural fluid of TB and non-TB patients determined by SCENITH. Representative flow cytometry histograms show puromycin incorporation (anti-puromycin MFI) after treatment with vehicle (C, Control), 2-deoxy-D-glucose (DG), oligomycin (O), or both inhibitors (DG+O). **(B)** Metabolic profiling of unstimulated (M0) or LPS/IFNγ-activated (M1) macrophages treated or not with RvD5 (33 nM) determined by SCENITH. Representative histograms of puromycin incorporation are shown. **(C)** Quantification of anti-puromycin MFI values (n=4). Arrows and adjacent numbers indicate the differences in MFI used to calculate metabolic parameters: (1) glucose dependence, (2) mitochondrial dependence, (3) glycolytic capacity, and (4) fatty acid and amino acid oxidation capacity (FAO/AAO). Bar graphs show glucose dependence and FAO/AAO capacity for each condition. Data was analyzed by one-way ANOVA with

Dunnett's multiple comparisons test. All quantitative data are presented as scatter plots of individual donors, with means ± SEM.**Figure S5. RvD5 does not induce a foamy phenotype in macrophages.** Human monocyte-derived macrophages were treated for 24 h with 20% v/v TB pleural effusion (PE), RvD5 (33 nM), or vehicle (ethanol), followed by staining with Oil Red O to assess lipid accumulation. **(A)** Quantification of lipid content based on the raw integrated density of Oil Red O staining per field. **(B)** Representative bright-field microscopy images of macrophages stained with Oil Red O (40× magnification). Statistical analysis: Nested t-test from two independent experiments, with 5 images analyzed per condition. condition.

**Figure S6 RvD5 Impairs Glycolysis Without Altering Mitochondrial Bioenergetics in M1 macrophages. (A)** Metabolic phenogram plotting basal oxygen consumption rate (OCR) versus basal extracellular acidification rate (ECAR) in unstimulated (M0) or LPS/IFNγ-activated (M1) macrophages, treated or not with RvD5 (33 nM), n=6. **(B, C)** Representative traces from a single experiment showing **(B)** OCR and **(C)** proton efflux rate (PER) for ATP rate determinations in M0, M1, and RvD5-treated M1 macrophages (means ± SEM of triplicates shown). **(D)** ATP production rates from mitochondrial oxidative phosphorylation (MitoATP) and glycolysis (GlycoATP) in M0 and M1 macrophages, treated or not with RvD5 (n=5). Data was analyzed by one-way ANOVA with Dunnett's multiple comparisons test. All quantitative data are presented as scatter plots of individual donors, with means ± SEM shown. Data obtained from Seahorse experiments were normalized based on the area covered by the cells, and a scale factor of 10,000 was used.

**Figure S7. RvD5 Does Not Alter Mitochondrial Mass or Morphology in M1 macrophages. (A)** Mitochondrial mass measured by mean fluorescence intensity (MFI) of a Mitospy probe in unstimulated (M0) or LPS/IFNγ-activated (M1) macrophages, with or without RvD5 (n=8). **(B)** Representative histogram overlays. **(C)** Mitochondrial morphology assessed by transmission electron microscopy (TEM). Left panels show representative TEM micrographs of M0, M1, and RvD5-treated M1 macrophages, with mitochondria pseudocolored in lilac. Right panels show morphometric analysis of mitochondria based on 16 images of single cells from two independent donors per condition. Mitochondrial mass data (B) were analyzed by one-way ANOVA followed by Sidak's multiple comparisons test. Mitochondrial morphology data (C) were analyzed using a mixed-effects model followed by Dunnett's multiple comparisons test.

**Figure S8. RvD5 Reduces Glycolysis Through GPR32 receptor.** Metabolic phenogram plotting basal oxygen consumption rate (OCR) versus basal extracellular acidification rate (ECAR) in RvD5-treated M1 macrophages, with or without neutralizing anti-GPR32 antibody (αGPR32) (n=3). **(B, C)** Representative traces from a single experiment showing **(B)** OCR and **(C)** proton efflux rate (PER) for ATP rate determinations in RvD5-treated M1 macrophages, with or without αGPR32. Data are presented as means ± SEM of triplicates. **(D)** ATP production rates from mitochondrial oxidative phosphorylation (MitoATP) and glycolysis (GlycoATP) in RvD5-treated M1 macrophages, with or without αGPR32 (n=3). Data was analyzed by paired t-test. All quantitative data are presented as scatter plots of individual donors, with means ± SEM.

**Figure S9. RvD5 Promotes Pro-Resolving Functions in M1 Macrophages. (A)** Neutrophil efferocytosis. Apoptosis of human neutrophils was induced and confirmed via

Annexin V/propidium iodide staining; apoptotic cells were labeled with CFSE. FSC vs. SSC plots show gating of isolated apoptotic neutrophils and macrophages. CFSE-labeled apoptotic neutrophils were co-cultured with unstimulated or LPS/IFNγ-activated (M1) macrophages in the presence or absence of RvD5. The percentage of macrophages that ingested $CFSE^+$ apoptotic neutrophils was quantified (n=7). **(B)** Surface expression of MerTK on M0, M1, and RvD5-treated M1 macrophages (n=6). Data was analyzed by one-way ANOVA with Dunnett's multiple comparisons test. **(C)** IL-10 secretion measured by ELISA in supernatants from M0, M1, and RvD5-treated M1 macrophages after 24 h (n=8). Data were analyzed by the Friedman test with the two-stage linear step-up procedure of Benjamini, Krieger and Yekutieli. **(D)** Surface expression of CD36 and CD14. Mean fluorescence intensity (MFI) of CD36 and CD14 on unstimulated (M0) or LPS/IFNγ-activated (M1) macrophages, with or without RvD5 (n=3). Data was analyzed by Friedman test with Dunn's multiple comparisons test. All quantitative data are presented as scatter plots of individual donors, with means ± SEM.

**Figure S10. Antigen-Presenting Cell-Dependent Responses Are Impaired by RvD5. (A-D)** Peripheral blood mononuclear cells (PBMC) were stimulated with irradiated Mtb (iMtb) in the presence of RvD5 or vehicle (ethanol) for six days. **(A)** Percentage of $CD4^+$ T cells expressing the activation marker CD69 (n=6). **(B)** Representative flow cytometry dot plots showing CD4 versus CD69 expression from one experiment. **(C)** Percentages of $CD4^+$ T cells expressing CD25 (n=6). **(D)** Representative dot plots showing CD4 versus CD25 expression from one experiment. **(E-G)** PBMC were stimulated with antibodies against CD28 and CD3 in the presence of RvD5 or vehicle (ethanol) for two days. **(E)** Percentage of $CD4^+$ T cells expressing CD69 (n=6). **(F)** Percentage of $CD4^+$ T cells expressing CD25 (n=6). **(G)** Representative dot plots from one experiment showing CD4, CD69, and CD25 expression. Data were analyzed by two-way ANOVA followed by Sidak's multiple comparisons test. **(H-I)** PBMCs were exposed to RvD5 in the presence or absence of rapamycin (Rap, 50 nM), an autophagy activator, and then stimulated with iMtb. The graphs show the percentage of $CD4^+$ T cells expressing the activation markers CD69 **(H)** and CD25 **(I)** (n = 4 donors per group). Statistical significance was determined by two-way ANOVA followed by the two-stage linear step-up procedure of Benjamini, Krieger and Yekutieli for FDR correction across all pairwise comparisons. Data are presented as scatter plots of individual donors, with means ± SEM.

**Figure S11. HIF1α activation Restores RvD5-Induced Pro-Resolving Functions in M1 Macrophages. (A)** Unstimulated (M0) or LPS/IFNγ activated (M1) macrophages were treated with RvD5 in the presence or absence of the HIF 1α stabilizer DMOG. Co cultures with CFSE labeled apoptotic neutrophils were analyzed by flow cytometry to determine the percentage of macrophages that ingested apoptotic neutrophils (n=6). **(B)** MerTK surface expression on M0 and M1 macrophages treated with RvD5 and/or DMOG (n=7). **(C)** IL 10 secretion measured by ELISA in supernatants from M0 and M1 macrophages treated with RvD5 and/or DMOG for 24 h (n=8). All data were analyzed by one way ANOVA followed by Sidak's multiple comparisons test. Data are presented as scatter plots of individual donors, with means ± SEM.
